# *C. elegans* hermaphrodites undergo semelparous reproductive death

**DOI:** 10.1101/2020.11.16.384255

**Authors:** Carina C. Kern, Shivangi Srivastava, Marina Ezcurra, Nancy Hui, StJohn Townsend, Dominik Maczik, Victoria Tse, Jürg Bähler, David Gems

**Author notes:** **Corresponding author:** David Gems, Institute of Healthy Ageing, and Department of Genetics, Evolution and Environment, University College London, Gower Street, London WC1E 6BT, United Kingdom.

## Abstract

Ageing in the nematode *Caenorhabditis elegans* is unusual in terms of the severity and early onset of senescent pathology, particularly affecting organs involved in reproduction (Ezcurra et al., 2018; Garigan et al., 2002; Herndon et al., 2002). In post-reproductive *C. elegans* hermaphrodites, intestinal biomass is converted into yolk leading to intestinal atrophy and yolk steatosis (Ezcurra et al., 2018; Sornda et al., 2019). We recently showed that post-reproductive mothers vent yolk which functions as a milk (*yolk milk*), supporting larval growth that is consumed by larvae (Kern et al., 2020). This form of massive reproductive effort involving biomass repurposing leading to organ degeneration is characteristic of semelparous organisms (i.e. that exhibit only a single reproductive episode) ranging from monocarpic plants to Pacific salmon where it leads to rapid death (reproductive death) (Finch, 1990; Gems et al., 2020). Removal of the germline greatly increases lifespan in both *C. elegans* and Pacific salmon, in the latter case by suppressing semelparous reproductive death (Hsin and Kenyon, 1999; Robertson, 1961). Here we present evidence that reproductive death occurs in *C. elegans*, and that it is suppressed by germline removal, leading to extension of lifespan. Comparing three *Caenorhabditis* sibling species pairs with hermaphrodites and females, we show that lactation and massive early pathology only occurs in the former. In each case, hermaphrodites are shorter lived and only in hermaphrodites does germline removal markedly increase lifespan. Semelparous reproductive death has previously been viewed as distinct from ageing; however, drawing on recent theories of ageing (Blagosklonny, 2006; de Magalhães and Church, 2005; Maklakov and Chapman, 2019), we argue that it involves exaggerated versions of programmatic mechanisms that to a smaller extent contribute to ageing in non-semelparous species. Thus, despite the presence of reproductive death, mechanisms of ageing in *C. elegans* are informative about ageing in general.

---

In this study, using a comparative biology approach, we explore the hypothesis that *C. elegans* experience semelparous reproductive death (Gems et al., 2020; Kern et al., 2020). Protandrous hermaphroditism (where sperm are produced first and then oocytes) allows *C. elegans* to rapidly colonize new food patches, but at the cost of a very short reproductive span due to sperm depletion (Hodgkin and Barnes, 1991; Schulenburg and Félix, 2017). The capacity to convert somatic biomass into yolk milk to feed to offspring allows post-reproductive *C. elegans* to reduce this cost, and promote inclusive fitness (Kern et al., 2020). This predicts that, among species of *Caenorhabditis*, lactation will occur in androdioecious (hermaphrodite, male) but not gonochoristic (female, male) species. To test this, we compared three pairs of sibling species in this genus, where one is androdioecious and the other gonochoristic: *C. elegans* vs *C. inopinata*, *C. briggsae* vs *C. nigoni*, and *C. tropicalis* vs *C. wallacei* (Fig. 1a). These pairs represent three independent occurrences of the evolution of hermaphroditism from gonochoristic ancestors (Kiontke et al., 2011). For each pair, both yolk venting and copious laying of unfertilised oocytes (which act as vectors for yolk milk (Kern et al., 2020)) was seen in hermaphrodites but not females (Fig. 1b,c; Extended data Fig. 1a,b). As in *C. elegans*, vitellogenin (yolk protein) accumulation continued into later life to high levels in *C. briggsae* and *C. tropicalis* hermaphrodites, but not in *C. inopinata*, *C. nigoni* and *C. wallacei* females (Extended Data Fig. 1c,d). We also compared an androdioecious-gonochoristic sibling species pair from the free-living nematode genus *Pristionchus* and, again, only the former vented yolk milk and oocytes, and accumulated vitellogenin internally in later life (Fig. 1b,c, Extended Data Fig. 1a-d). *Pristionchus* proved to lack the larger vitellogenin species corresponding to YP170 in *Caenorhabditis*; instead accumulation of only the smaller species equivalent to *Caenorhabditis* YP115/YP88 was seen (Extended Data Fig. 1c,d).

**Fig. 1.**
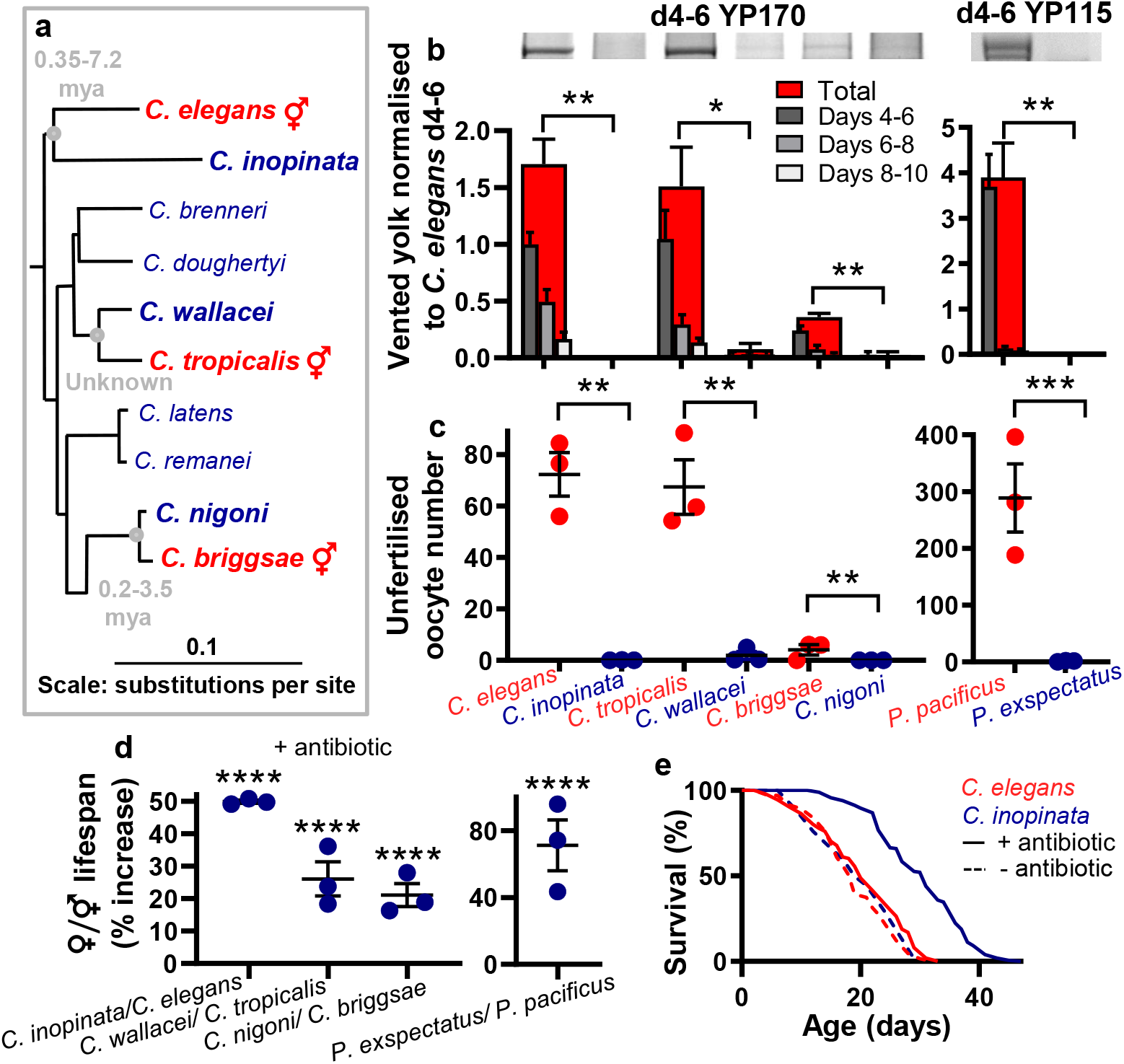
*Caenorhabditis* hermaphrodites exhibit yolk milk venting and a shorter lifespan. **a**, Phylogenetic tree showing androdioecious and gonochoristic *Caenorhabditis* sibling species (Stevens et al., 2019). Estimates of time since divergence: *C. elegans, C. inopinata* 0.35-7.2 MYA; *C. briggsae, C. nigoni* 0.2-3.5 MYA (Cutter et al., 2019); *C. tropicalis, C. wallacei* still unknown. **b**, Yolk milk venting present in hermaphrodites but not in females (n=100 per trial). Top: major YP at the peak of venting. Bottom: Quantitative data normalised to total yolk on d4-6 in *C. elegans*. **c**, Unfertilised oocytes are laid by hermaphrodites but not females (n=10 per trial). **d**, Females are longer lived than sibling species hermaphrodites (unmated, with antibiotic). **b**,**c**,**d**, Mean ± s.e.m. of 3 trials. **b**,**c**, One-way ANOVA or unpaired t-test. **d**, log rank test. * *p*<0.05, ** *p*<0.01, *** *p*<0.0001, **** *p*<0.00001. **e**, *C. inopinata* is longer-lived than *C. elegans* when bacterial infection is prevented using an antibiotic (carbenicillin) (this figure) or UV irradiation of *E. coli* (Extended data Fig. 2d). For statistical details, see Extended Data Table 1, and for raw lifespan data Supplementary File 1.

Based on our semelparity hypothesis, the presence of lactation in hermaphrodites but not females predicts that only the former will exhibit reproductive death, i.e. that females should be longer lived and show greatly reduced pathology. Consistent with this, lifespan was longer in females than in hermaphrodites for the *C. tropicalis*/*C. wallacei* and *C. briggsae*/*C. nigoni* pairs but, as previously reported (Woodruff et al., 2018), this was not the case for the *C. elegans*/*C. inopinata* pair (Extended Data Fig. 2a,b). To exclude possible species differences in susceptibility to infection by the *E. coli* food source (Gems and Riddle, 2000), lifespan was measured in the presence of antibiotics, and here females were longer lived in all three sibling species pairs (Fig. 1d, e, Extended Data Fig. 2a,b); this result suggests that *C. inopinata* are hyper-susceptible to bacterial infection, perhaps reflecting their distinct natural environment (syconia on Okinawan fig trees). The optimal culture temperature for *C. inopinata* is higher (25-29°C) than that for *C. elegans* (20-22°C) (Kanzaki et al., 2018), raising the possibility that culture of *C. inopinata* at the sub-optimally low temperature of 20°C might cause a disproportionately low rate of living, thereby increasing lifespan relative to *C. elegans*; however, *C. inopinata* was also longer lived than *C. elegans* at 25°C (Extended Data Fig. 2c), arguing against this possibility. Similarly, in the *Pristionchus* sibling species pair, the hermaphrodites were shorter lived (Fig. 1d, Extended Data Fig. 2a,b), consistent with a previous report of greater longevity in *Pristionchus* females (Weadick and Sommer, 2016).

**Fig. 2.**
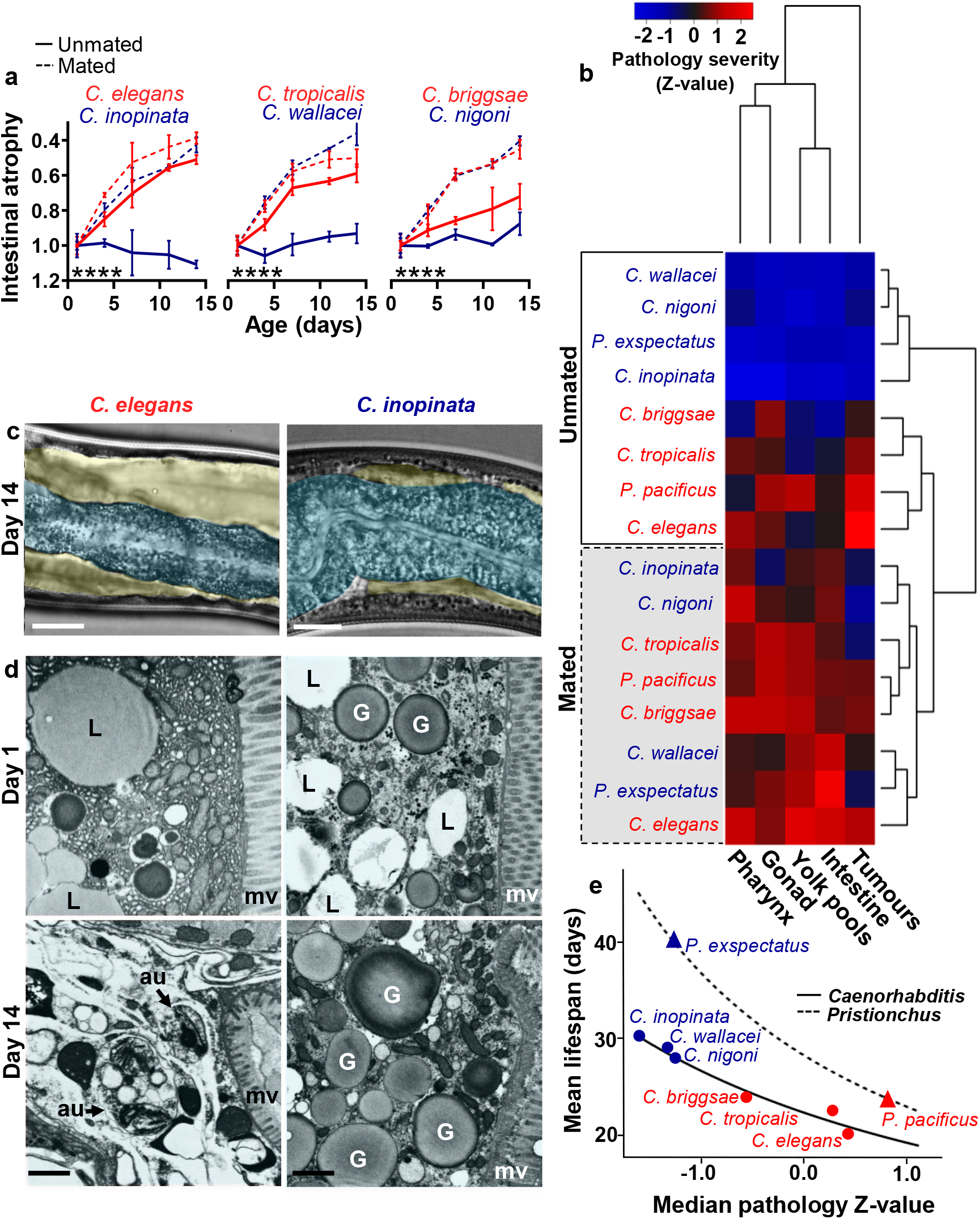
Reproductive death is constitutive in hermaphrodites and facultative in females. **a**, Intestinal atrophy is constitutive in hermaphrodites and mating-induced in females. Pathology score was normalised to d1 values for each species/treatment. Mean ± s.e.m. of 3 trials (n=10 per trial per time point); **** *p*<0.00001, ANCOVA. For measurement of pathologies, statistical comparisons between species and treatments, and representative images of pathologies used for scoring, see Extended Data Fig. 4. **b**, Reproductive death-associated pathologies are constitutive in hermaphrodites and induced in females by mating. The heat map compares differences in pathology progression across species and treatments by transforming calculated gradients from pathology measurements through time into Z-scores (c.f. Extended data Fig. 4a,d; n=10 per time point, with 5 time points up to d14, when pathology peaks for *C. elegans*). This allows comparison between different pathologies by normalising levels of a given pathology to the average level of that pathology in the group (represented by a Z-score of 0, with a Z-score of +2 representing the maximum pathology severity in the group and −2 the healthiest animals). Hierarchical clustering is based on pair-wise Euclidean distances. **c**, Intestinal atrophy (blue) and yolk pool accumulation (yellow) in ageing *C. elegans* but not *C. inopinata* (d14, unmated). Representative Nomarski images. Scale 25 μm. **d**, Severe degeneration of intestinal ultrastructure in ageing *C. elegans* but not *C. inopinata* (day 14, unmated). Note loss of organelles and ground substance. Representative TEM cross-section images through the intestine. mv, microvilli; au, autophagosome-like structure, L, lipid droplets; G, yolk or gut granules (tentative identification). Scale 1 μm (20,000x). For mated *C. inopinata* showing intestinal degeneration on d14, see Extended Data Fig. 5d,e. **e**, Correspondence between pathology severity and lifespan. Linear regression of the median of all pathology Z-scores (c.f. Fig. 2b) with lifespan. Y axis shows mean lifespan of each species (antibiotic treated). A line of best fit is drawn to connect points for the *Caenorhabditis* species; based on this a hypothetical line is drawn for the *Pristionchus* species. For regression of individual pathologies against lifespan, see Extended Data Fig. 2e.

Next we compared patterns of senescent pathology in the three sibling *Caenorhabditis* species pairs using Nomarski microscopy and transmission electron microscopy. Early, severe senescent pathologies in *C. elegans* adult hermaphrodites include prominent uterine tumours, gonadal atrophy and fragmentation, pharyngeal deterioration, as well as intestinal atrophy coupled to yolk steatosis (de la Guardia et al., 2016; Ezcurra et al., 2018; Garigan et al., 2002; Herndon et al., 2002; Wang et al., 2018). Senescent pathologies seen in *C. elegans* were also seen in the other two hermaphroditic species, but were largely absent from females, and this was true also of the *Pristionchus* sibling species pair (Fig. 2a,b,c). Striking degeneration of intestinal ultrastructure was seen in older *Caenorhabditis* hermaphrodites but not in females, consistent with earlier observations of *C. elegans* (Herndon et al., 2002; Herndon et al., 2018) and *C. briggsae* (Epstein et al., 1972), including prominent autophagosomes (Fig. 2d, Extended Data Fig. 3). Thus, females are longer lived and free of the severe pathology present in hermaphrodites.

The severity of early senescent pathology in hermaphrodites varied between species, with the ranking *C. elegans* > *C. tropicalis* > *C. briggsae* (Fig. 2a,b), and hermaphrodite longevity showed the inverse ranking (Extended data Table 1). Notably, the graded variation of pathology level with lifespan implies a continuum between presence and absence of reproductive death (i.e. between semelparity and iteroparity; see below) (Fig. 2e). This suggests that *C. elegans* and *C. briggsae* represent late and early stages, respectively, in the evolution of reproductive death. Notably, *C. briggsae*/*C. nigoni* inter-species mating can produce offspring (Baird et al., 1992), implying relatively recent divergence of these two species from a common ancestor.

The above comparisons were performed using unmated animals, meaning that hermaphrodites produced progeny but females did not. Thus, the differences observed could reflect the presence/absence of progeny production. To assess this hypothesis, we compared lifespan and senescent pathology in mated animals of the different species. Mating shortened lifespan in most species, as previously seen in *C. elegans* (Gems and Riddle, 1996; Shi and Murphy, 2014), and abrogated the greater longevity of females (Extended Data Fig. 5a-c). Mating also induced intestinal atrophy in females and enhanced it in hermaphrodites, and mated animals of all six species showed similar levels of intestinal atrophy (Fig. 2a). A similar effect of mating was also seen on a range of other pathologies, where similar levels of deterioration were seen in mated females and hermaphrodites, as shown by cluster analysis of quantified severity in pathology (Fig. 2b). These findings suggest that reproductive death is induced by mating in females. This provides a potential explanation for how such similar patterns of pathogenesis evolved independently in the three androdioecious species: as the consequence of a switch from facultative (mating-induced) reproductive death in females to constitutive reproductive death in hermaphrodites.

According to this model, the evolution of reproductive death in hermaphrodites requires only its activation in the absence of mating. Evolution of hermaphroditism in *C. elegans* likely began when females developed the capacity to generate and activate self sperm (Baldi et al., 2009). One possibility is that the emergence of self sperm was sufficient for reproductive death to become constitutive. However, spermless *fog-2(q71)* and *fem-2(e2006)* female mutant *C. elegans* showed no detectable reduction in intestinal atrophy (Extended Data Fig. 5f), arguing against this hypothesis. This result and the less severe senescent pathology seen in *C. briggsae* suggest that the evolution of constitutive reproductive death is a complex, multi-step process.

In semelparous species where reproductive death is triggered upon reproductive maturity, pre-empting reproduction can greatly increase lifespan (Finch, 1990; Gems et al., 2020). For example, removal of flowers prior to pollination can increase lifespan in soybean from 119 to 179 days (Leopold et al., 1959), and gonadectomy of Pacific salmon before spawning can increase maximum lifespan from 4 years to up to 8 years (Robertson, 1961). Similarly, prevention of germline development in *C. elegans* hermaphrodites greatly increases lifespan (Hsin and Kenyon, 1999) and suppresses intestinal atrophy (Ezcurra et al., 2018). One possibility is that germline ablation extends *C. elegans* lifespan by suppressing reproductive death. Given the absence of reproductive death in unmated *Caenorhabditis* and *Pristionchus* females, this predicts that germline ablation will increase lifespan more in hermaphrodites than in females, and this proved to be the case. Germline ablation using laser microsurgery (Hsin and Kenyon, 1999) suppressed major pathologies and caused large increases in lifespan in hermaphrodites, but only marginal increases in lifespan in females (Fig. 3a,b). We also surveyed published reports that have assessed effects on lifespan of gonadectomy or, for some semelparous species, prevention of reproductive death by other means. This confirmed that large increases in lifespan are typical of semelparous but not iteroparous organisms (Fig. 3b). Lifespans of germline-ablated animals in each sibling species pair were similar (Extended Data Fig. 6), supporting the view that the shorter lifespan of hermaphrodites is attributable to reproductive death.

**Fig. 3.**
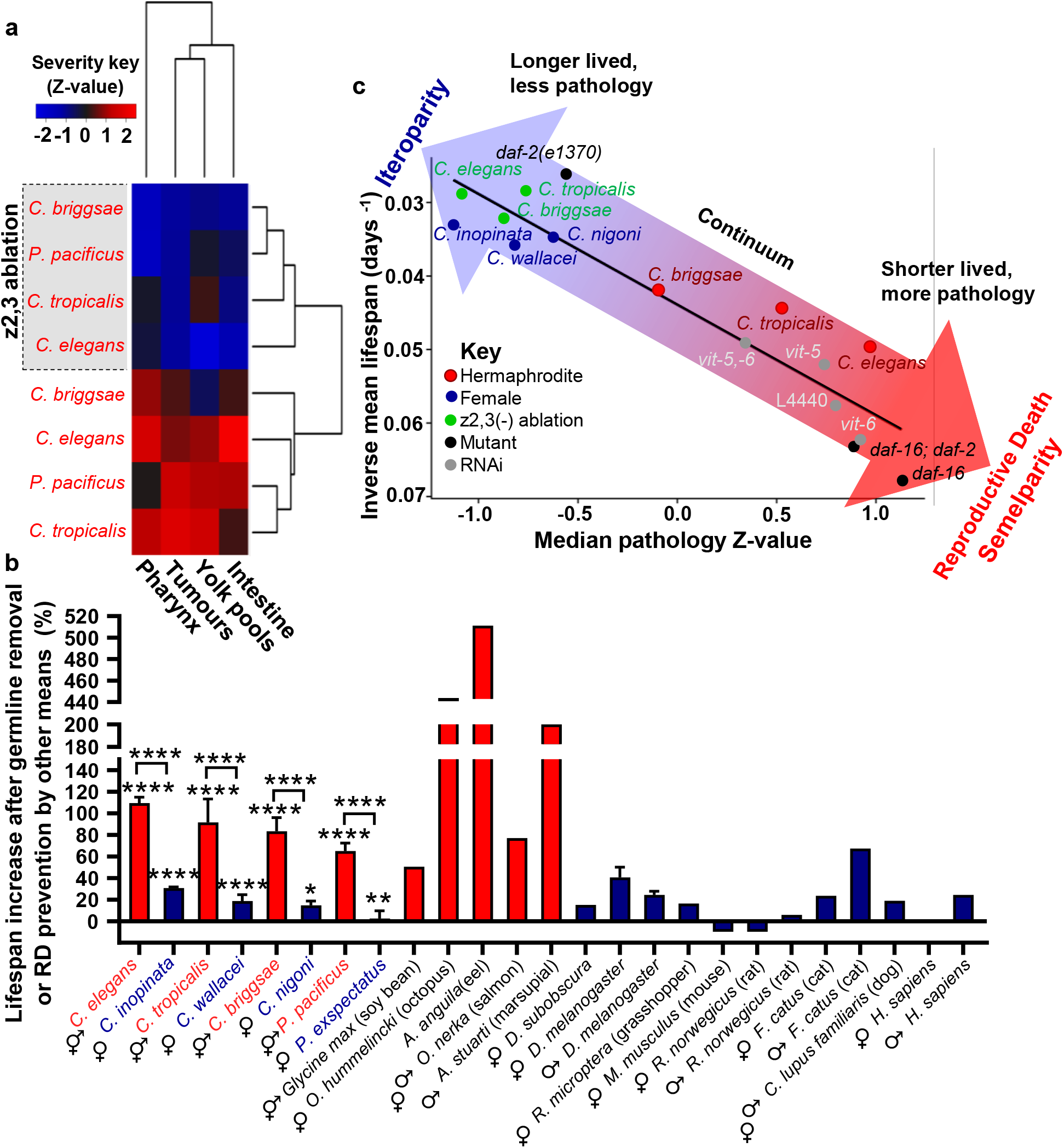
Germline ablation extends hermaphrodite lifespan by suppressing reproductive death. **a**, Germline ablation strongly suppresses pathology progression in hermaphrodites (c.f. Fig. 2b; n=5-10 per time point). For full pathology measurements, see Extended data Fig. 6a; for pathology progression in germline-deficient mutants as well as representative images used to score pathologies, see Extended Data Fig. 6b,c. **b**, Germline ablation strongly increases lifespan in hermaphrodites but not female nematode species. Similarly, blocking reproductive death in semelparous species strongly increases lifespan, while gonadectomy in iteroparous species overall does not. For *Caenorhabditis* and *Pristionchus* species, mean ± s.e.m. of 3 trials is shown (stars, log rank test for ablated *vs* control animals, and Cox proportional hazard test to compare effect in hermaphrodites *vs* females). Values for other species are taken from a recent review (Gems et al., 2020); for multiple studies of the same species, only references with both male and female data are included, and for *A. anguila* (eel) data from the most recent source (Tesch, 1977). For individual trials, details on statistics and raw lifespan data, see Extended Data Fig. 7, Extended data Table 2 and Supplementary File 1, respectively. **c**, Correlation between reproductive death-associated pathologies and lifespan in different species and following different treatments in *C. elegans*, supporting the presence of a semelparity-iteroparity continuum. X axis, median pathology Z-score. Y axis, mean lifespan (unmated, antibiotic treated). Other data sources: lifespan and pathology with RNAi lifespan (Sornda et al., 2019); lifespan of mutants (Bansal et al., 2014). For mutant pathology measurements, see Extended data Fig. 8.

Results presented in this study, taken together with earlier findings, show that properties of *C. elegans* fit the criteria used to define reproductive death in semelparous species, i.e. that it exhibits reproductive death. *C. elegans* develop massive, severe pathology in organs linked to reproduction relatively early in adulthood, some of which result from biomass repurposing for yolk milk production (Ezcurra et al., 2018; Gems et al., 2020; Kern et al., 2020), while none of this is seen in unmated females. Hermaphrodites are shorter lived than females of their sister species, but not after germline removal, which causes large increases in lifespan only in hermaphrodites. This is consistent with the view that germline removal, like gonadectomy in Pacific salmon, increases lifespan by preventing reproductive death. Consistent with this, *C. elegans* males resemble *Caenorhabditis* females in not exhibiting major degenerative pathologies such as intestinal atrophy and gonad disintegration (de la Guardia et al., 2016; Ezcurra et al., 2018), and germline removal in wild-type *C. elegans* males has little effect on lifespan (McCulloch, 2003). Moreover, prevention of reproductive maturity by gonadectomy or other means typically causes large increases in lifespan in semelparous organisms but not iteroparous organisms (with multiple rounds of reproduction) (Gems et al., 2020) (Fig. 2b).

The occurrence of reproductive death in *C. elegans* has profound implications both in terms of understanding the biology of ageing, and what one can learn from *C. elegans* as a model for ageing in general. Insofar as senescent pathologies develop as a cost of lactation (Kern et al., 2020), they are not the result of futile run-on of programmes that promoted fitness earlier in life (or quasi-programmes (Blagosklonny, 2006)), as was previously suggested (Ezcurra et al., 2018; Herndon et al., 2002; Sornda et al., 2019). However, fitness-promoting processes to which uterine tumour formation is coupled have not (yet) been identified; thus, like the ovarian teratomas that they resemble (Wang et al., 2018), uterine tumours do appear to the be result of quasi-programmes. Thus, both costly programmes (Gems et al., 2020) and quasi-programmes are operative in *C. elegans* hermaphrodite senescence.

The most exciting thing about the discovery of interventions producing large increases in lifespan in *C. elegans* was the implied existence of mechanisms with powerful effects on ageing as a whole. Combined with the notion that ageing rate was a function of somatic maintenance (Kirkwood and Rose, 1991), this suggested that similar plasticity might exist in humans, affecting the entire ageing process, and providing a target for future interventions to greatly decelerate ageing. Here we present evidence that lifespan in *C. elegans* is limited by reproductive death; we have also argued elsewhere that reproductive death is permissive for the evolution of the rare phenomenon of programmed adaptive death, which further shortens lifespan, and that this has happened in *C. elegans* (Galimov and Gems, 2020a, b; Lohr et al., 2019). Suppression of such programmatic etiologies of ageing provides a potential explanation for the unusually large magnitude of increases in lifespan seen in *C. elegans*. This raises the possibility that in iteroparous organisms no core mechanisms of ageing as a whole exist that are amenable to manipulation to produce dramatic deceleration of ageing. Sadly, our findings in certain respects explain away the mystique of *C. elegans* ageing.

Does the occurrence of reproductive death mean that *C. elegans* die from mechanisms unrelated to those operative in most other organisms? We believe not. In the past, a sharp distinction was drawn between true ageing, caused by stochastic damage accumulation, and programmed ageing as observed in plants (e.g. leaf senescence) and semelparous species like Pacific salmon (Finch, 1990; Gems et al., 2020). However, it has been recognized that evolutionarily conserved regulators of phenotypic plasticity in ageing, such as the insulin/IGF-1 and mechanistic target of rapamycin (mTOR) pathways, can act through programmatic mechanisms to promote senescent pathology (Blagosklonny, 2006; de Magalhães and Church, 2005; Gems and de la Guardia, 2013; Maklakov and Chapman, 2019). Inhibiting these pathways can cause large increases in *C. elegans* lifespan but also smaller effects in higher animals; for example, mutation of phosphatidylinositol 3-kinase can increase median lifespan by up to ~10-fold in *C. elegans,* but only ~1.07-fold and ~1.02-fold in *Drosophila* and mouse, respectively (Ayyadevara et al., 2008; Foukas et al., 2013; Slack et al., 2011). Given the evolutionary conservation of the role of these pathways in ageing, we suggest that mechanisms of reproductive death evolve by repurposing of programmatic mechanisms that affect ageing in iteroparous organisms (Gems et al., 2020). It has been argued that semelparous and iteroparous life histories are not isolated phenomena but rather represent a continuum (Hughes, 2017). Supporting this is the observed gradient between the presence and absence of reproductive death across nematode species, and in *C. elegans* across interventions that suppress pathologies of reproductive death and extend lifespan, including germline ablation and reduced IIS (Fig. 3c). Thus, *C. elegans* is a good model for understanding programmatic mechanisms of ageing that lead to senescent multimorbidity that are universal across metazoan organisms.

## Supporting information

Supplementary Guide

## Acknowledgments

We are grateful to V. Konstantellos for technical assistance, M. Turmaine for help with electron microscopy, A. Barrios and R.J. Poole for access to laser ablation facilities, N. Kanzaki and T. Kikuchi (F.F.P.R.I. Tsukuba) for providing *C. inopinata*, and R.J. Sommer (M.P.I. Developmental Biology, Tübingen) for providing *Pristionchus* species. We also thank A.D. Cutter (University of Toronto), M.-Q. Dong and C. Zhai (N.I.B.S. Beijing), D. Hall (Albert Einstein College of Medicine), R.J. Sommer (Tübingen), V. Rottiers (U.C. Berkeley) and E.R. Galimov, Y. de la Guardia and other members of the Gems lab for advice and useful discussion and/or comments on the manuscript. Some strains were provided by the Caenorhabditis Genetics Center, which is funded by NIH Office of Research Infrastructure Programs (P40 OD010440). S.T. was supported by a Boehringer Ingelheim Fonds PhD Fellowship. This work was supported by a Wellcome Trust Strategic Award (098565/Z/12/Z) and a Wellcome Trust Investigator Award (215574/Z/19/Z) to D.G..

## Author contributions

D.G. supervised the project. C.C.K. and D.G. conceived the project, designed the experiments and data analysis, and wrote the manuscript. C.C.K., N.H., M.E., S.S. and V.T. performed experiments. D.M., N.H., M.E. and V.T. performed the pathology data capture and analysis. N.H. assisted with the collection and measurement of internal and vented proteins. S.S. scored lifespans. S.S. and N.H. performed laser ablations. S.T. and J.B. performed and interpreted quantitative analysis of pathology and comparison to lifespan.

