## Extended Data and Methods for "*C. elegans* hermaphrodites undergo semelparous reproductive death"

### Supplementary Information Guide

<u>**This file**</u>

**Methods.**

**Extended Data Figs. 1-8.**

**Extended Tables 1-3.**

<u>**Additional files**</u>

**Supplementary File 1 | Raw lifespan data.** This contains the raw data from which survival curves and calculated lifespan values were derived. Raw data is provided in Zeihm tables, which apply minimal reporting standards to ensure inclusion of all essential information about experimental conditions. Zeihm tables allow future reanalysis of survival data in published studies.

#### Methods

No statistical methods were used to predetermine sample size. The experiments were not randomised. The investigators were not blinded to allocation during experiments and outcome assessment unless otherwise stated. All statistical tests were performed on raw data using GraphPad Prism 8.0 unless otherwise stated.

##### Nematode culture methods and strains

Maintenance of *C. elegans* and other species was performed using standard protocols (Brenner, 1974). Unless otherwise stated, all strains and species were grown at 20˚C on nematode growth media (NGM) with plates seeded with *E. coli* OP50 to provide a food source. For *C. elegans,* a N2 hermaphrodite stock recently obtained from the Caenorhabditis Genetics Center was used as wild type (N2H) (Ref. (Zhao et al., 2019)). Genotypes of *C. elegans* mutant strains used are described in Wormbase (www.wormbase.org), and included CB3844 *fem-3(e2006)*, CB4037 *glp-1(e2141),* GA14 *fog-2(q71)*, GA114 *daf-16(mgDf50)*; *daf-2(e1370),* GA1928 *daf-2(e1370)*, GR1307 *daf-16(mgDf50)* and SS104 *glp-4(bn2)*. Other nematode species used were *C. inopinata* (NK74SC), *C. remanei* (SB146), *C. tropicalis* (JU1373), *C. wallacei* (JU1873), *C. briggsae* (VT847), *C. nigoni* (JU1325), *P. pacificus* (RS2333) and *P. exspectatus* (RS5522). *C. inopinata* were raised at 25˚C until the L4 stage and then transferred to 20˚C since at the latter temperature they exhibit a very slow development rate(Kanzaki et al., 2018). Temperature sensitive mutants (*fem-3, fog-2 glp-1*, *glp-4*) and accompanying controls were raised at 15˚C until the L4 stage and then shifted to 25˚C, the non-permissive temperature.

##### Nomarski microscopy

Live nematodes were placed onto 2% agar pads and anaesthetised in a drop of 0.2% levamisole, with cover slips gently placed on top. Images were captured using either a Zeiss Axioskop 2 plus microscope with a Hamamatsu ORCA-ER digital camera C4742-95 and Volocity 6.3 software (Macintosh version) for image acquisition; or an ApoTome.2 Zeiss microscope with a Hamamatsu digital camera C13440 ORCA-Flash4.0 V3 and Zen software. Brightness and contrast were adjusted equally across the entire image, and where applicable applied equally to controls.

##### Transmission electron microscopy

Nematode sample preparation was based on a standard protocol(Shaham, 2006) (protocol 8). Briefly, animals were washed twice in M9 buffer, placed in fixing solution (2.5% glutaraldehyde, 1% paraformaldehyde in 0.1 M sucrose, 0.05 M cacodylate) and, to increase staining dye penetration, cut below the pharynx to remove the heads using a 30 G hypodermic needle. They were then rinsed 3 times in 0.2 M cacodylate, fixed in 0.5% $OsO_4$ and 0.5% $KFe(CN)_6$ in 0.1 M cacodylate on ice and sequentially washed in 0.1 M cacodylate and 0.1 M sodium acetate. Nematodes were then stained in 1% uranyl acetate in 0.1 M sodium acetate (pH 5.2) for 60 min, rinsed 3 times in 0.1 M sodium acetate and then rinsed overnight in Milli-Q water. Samples were then embedded in 3% seaplaque agarose, dehydrated and infiltrated using ethanol and propylene-resin series and then cured at 60˚C for 3 days. Serial 1 µm sections were taken for light microscopy, and, at the posterior half after the uterus, ultra-thin sections were cut at 70–80 nm using a diamond knife on a Reichert ultramicrotome. Comparative low magnification images showing halves of nematode sibling species pairs were taken from approximately the same region of the animals containing the posterior intestine. Sections were collected on formvar-coated 2x1 mm slot grids which allowed low magnification images to be taken without grid bars and stained with lead citrate before being viewed and imaged in a Joel 1010 Transition Electron Microscope (TEM). Images were recorded using a Gatan Orius camera, and obtained using Gatan imaging software. Brightness and contrast were adjusted equally across the entire image, and where applicable applied equally to controls.

**Lifespan measurements**

Animals were maintained at a density of 25-30 per plate from the egg stage to minimise population density-associated effects seen in *C. elegans* (Ludewig et al., 2017), and without the addition of FUDR. Nematode cohorts were scored every 2-3 days for the dead, and all animals transferred daily during the reproductive period, and every 6-7 days thereafter. Worms were scored as alive if they were motile or responded at all to gentle touch with a platinum worm-pick. For survival assays on UV-irradiated bacteria, 80 µl of *E. coli* OP50 was added to each NGM plate and left overnight at 20˚C. Plates were then exposed to UV for 30 min in a Stratalinker UV oven. Nematodes were raised on UV-treated bacteria from aseptic eggs. Raw mortality data for all trials are provided in Ziehm tables (Zhao et al., 2017) in Supplementary File 1 to enable close scrutiny of data (including future reanalysis). Survival plots show combined lifespan data and individual trials, with the L4 stage of development taken as day 0. Animals which disappeared from the plate, were accidently killed during handling, died due to desiccation on the Petri dish wall, ruptured through the vulva or died due to internal hatching of larvae were recorded as censored. Censored values were taken into account in statistical analysis. Tests for statistically significant differences between survival of cohorts employed the log rank test, and for significant differences in magnitude of effects of two treatments Cox proportional hazard analysis, in both cases performed using JMP software, version 14.0 (SAS Institute, Inc.).

**Counts of numbers unfertilized oocyte laid**

L4 larvae were maintained individually on 35 mm NGM plates with *E. coli* OP50 lawns (n = 10 per trial), and transferred every 24 hrs until unfertilized oocyte production ceased, with a minimum of 12 days scored for all worms. For scoring, the open Petri dish was place on an inverted plate bearing a grid of parallel lines, allowing the plate to be efficiently scored with a series of vertical sweeps. Where clumps of oocytes were seen, these were separated using a worm pick, to count individual oocytes. *P. pacificus* and *P. exspectatus* also laid some dead eggs, consistent with previous observations (R.J. Sommer, personal communication), which were discounted.

**Gel electrophoresis and quantitation of yolk proteins**

Quantitation of vented yolk protein was performed as previously described (Kern et al., 2020). Briefly, 100-200 L4 larvae were maintained on 35 mm plates with *E. coli* OP50 lawns and transferred daily to new plates. Following transfer, yolk milk was washed off with 1 ml of M9 containing 0.001% NP-40 to solubilise vitellogenin, as described (Sharrock, 1983), and 2 µg/ml BSA as an external standard. It was then centrifuged to separate out fractions prior to lyophilisation. After lyophilisation yolk was resuspended in a solution of 30 µl 4% SDS solution pH 9, 1 ml 1M Tris HCL pH 8, 5 ml Milli-Q water, 10 µl 0.5M EDTA, and 12 mg bromophenol blue. Quantitation of internal (non-vented) yolk used a protocol similar to that previously described (Sornda et al., 2019). Briefly, 10 animals per condition per treatment were transferred to NGM plates lacking bacteria and allowed to crawl for 30-60 sec to remove bacteria from the nematode surface. They were then transferred to 25 µl M9 and immediately frozen at -80˚C until used. Samples were mixed with 25 µL of 2× Laemmli sample buffer (Sigma-Aldrich), incubated at 70˚C and vortexed periodically for 15 min, and then incubated at 95˚C for 5 min and centrifuged at 6,000 rpm for 15 min. For all samples, sodium dodecyl sulfate–polyacrylamide gel electrophoresis (SDS-PAGE) was then performed, using Criterion XT Precast Gels 4–12% Bis-Tris (Invitrogen) and XT MOPS (Invitrogen) as a running buffer (7:1 ratio with Milli-Q water) at 90 V. Gels were stained with colloidal Coomassie blue as described (Kang et al., 2002), using 5% aluminum sulphate-(14-18)-hydrate and 2% orthophosphoric acid (85%) to create colloidal particles. Gels were analyzed using ImageQuant LAS 4000 (GE Healthcare). Protein band identification was based on published data (Depina et al., 2011).Within lanes, vented YPs were normalized to the BSA that had been added during sample collection as an external standard. For samples of internal YPs, YP bands were normalized to myosin as a standard to account for protein loss during gel loading, as well as to allow for normalisation to nematode size when comparing different species.

**Measurement of nematode senescent pathologies**

Nomarski microscopy images of senescent pathologies in cohorts of worms (n = 5-10 per time point per condition per trial) were captured on days 1, 4, 7, 11 and 14 and images analysed either quantitatively (intestine, yolk pools), or semi-quantitatively and blind (pharyngeal deterioration, gonad atrophy and fragmentation, uterine tumours) by several trained observers as described (Ezcurra et al., 2018; Garigan et al., 2002; Riesen et al., 2014). In brief, intestinal atrophy was quantified by measuring the intestinal width at a point between posterior gonad and anus, subtracting the width of the intestinal lumen, and dividing by the body width to obtain an estimate of intestinal cross-sectional width normalized to body size. Yolk pool accumulation was measured by dividing the area of yolk pools by the area of the body visible in the field of view at 630x magnification. For pharyngeal deterioration, gonad atrophy and fragmentation and uterine tumours, images were assigned scores of 1-5, where 1 = youthful, healthy appearance; 2 = subtle signs of deterioration; 3 = clearly discernible, mild pathology; 4 = well developed pathology; and 5 = tissue so deteriorated as to be barely recognizable (e.g., gonad completely disintegrated), or reaching a maximal level (e.g. large tumour filling the entire body width). Scoring was adapted to the different species being used as follows. Scoring of pharyngeal senescence in *Pristionchus* species took into account the normal absence of a grinder (Wei et al., 2003). For scoring of gonadal senescence, the criteria for a healthy gonad included whether the two arms of the gonad touch one another (which they do in young adults). Animals where the gonad arms do not touch, or which show narrowing on the gonad arms were given a score of 2, both features a score of 3, and in addition gonad arm fragmentation a score of 4. For uterine tumour scoring in young unmated females an empty uterus lacking eggs as was given a score of 1.

**Analysis and comparison of pathology progression**

In order to prepare data for modelling, data were pre-processed for the following reasons. Peak pathology level for *C. elegans* and most mated animals (which shows maximum pathology progression rate) is mostly reached by day 14, after which pathology levels plateau. Thus, pathology data after day 14 obscures calculation of gradients representing initial rate of pathology progression. Intestine scores for each combination of species and treatment were normalised to the respective mean intestine score on day 1. Therefore, normalised intestinal pathology scores represent the change in percentage of intestinal volume, accounting for differences in terms of the ratio of intestinal width to whole body width between species. For yolk pool scores no normalization was used because percentage of the body cavity containing yolk pools was measured. Yolk pools were analysed as score + 1, in order to ensure all data were positive (a requirement for Gamma regression). Generalised linear models (GLMs) were applied to the data to test the effect of species, treatment and trial on pathology progression. Interactions terms between these variables were also included. Different model families and link functions were systematically applied to each pathology dataset according to the type of data recorded. For positive, continuous variables (intestine and yolk pools), pathology scores were modelled as Gaussian and Gamma distributed variables. For ordinal variables (tumour, gonad and pharynx), cumulative link models were applied to the data using the *ordinal* package in R (Christensen, 2019). The following models were selected for each pathology based on minimisation of Akaine Information Criterion (AIC), meaning that these models explained the most variance in pathology progression using the fewest number of terms: (i) intestine: Gaussian distributed response variable with inverse link function; (ii) yolk pools: Gamma distributed response variable with identity link function; and (iii) tumour, gonad and pharynx: ordinal distributed response variable with log-gamma link function. The vast majority of trial terms were not statistically significant across all pathologies, indicating that differences in pathology progression were reproducible and robust. Hence, trial was not considered further as a variable. In order to compare whether differences in pathology progression between combinations of species and treatment were statistically significant, t-tests were applied explicitly to comparisons of interest using the *multcomp* package in R (Hothorn et al., 2008).

**Pathology relationship and comparison to lifespan**

In order to compare differences in pathology progression across species and treatments, the gradients from each model were transformed into Z-scores (which describe a value's relationship to the mean of a group of values), necessary to compare pathologies to one another. The pathology Z-scores are displayed and compared as heatmaps, using pairwise Euclidean differences to cluster pathologies and species/treatments according to profile similarity. In order to model the impact of pathology Z-scores on lifespan, linear regression was performed, using the pathology Z-score as the independent variable and the inverse of mean lifespan as the dependent variable. In order to assess the combined impact of all pathologies on lifespan, the median of the pathology Z-scores was used as the independent variable. The Z-scores for all pathologies were found to be statistically significant, and the median pathology Z-score was found to perform better than the individual pathologies.

**Ablation of the germline using laser microsurgery**

Live L1 larvae were mounted on a glass slide on a 5% agar pad with levamisole as anaesthetic. The concentration of levamisole and M9 buffer/Milli-Q water ratio was optimised for each sibling species pair: 0.2 mM levamisole in M9 for *C. elegans, C. inopinata, C. tropicalis* and *C. wallacei*; 0.2 mM levamisole in a 1:1 ratio of M9 to Milli-Q water for *C. briggsae* and *C. nigoni*; and 0.1 mM levamisole in M9 for *P. pacificus* and *P. exspectatus*. Ablations were performed using a Zeiss Axioplan 2 fitted with an Andor MicroPoint laser unit (CE N2 Laser with Ctlr/PSU/IntLk) at 440 nm and Dye Cell 435 nm filter. Germline precursor cells (Z2 and Z3) were identified by morphology and position with Nomarski optics and ablated in newly-hatched L1 animals using a standard protocol (Bargmann and Avery, 1995). Following ablation, larvae were transferred to fresh plates by washing them off with M9 buffer (30 μl) for all the pairs of species except for *C. briggsae* and *C. nigoni* which were recovered in a 1:1 ratio of M9 to Milli-Q water (total 30 μl). All ablated animals were then allowed to recover to day 1 of adulthood and checked under a Nikon SMZ645 microscope for both a lack of egg laying, indicative that the ablation was successful, as well as the presence of a vulva, indicating the Z1 and Z4 cells were intact. For unmated females which do not lay eggs, 15-20 randomly selected worms per condition per trial were checked under Nomarski (100x magnification with 10x air objective, without anaesthesia) on an NGM plate for the presence of a fully developed gonad, and none were found to have one. Mock treatment animals underwent the same manipulations as ablated animals with the exception of being shot with the laser microbeam.

Kanzaki, N., Tsai, I., Tanaka, R., Hunt, V., Liu, D., Tsuyama, K., Maeda, Y., Namai, S., Kumagai, R., Tracey, A., Holroyd, N., Doyle, S., Woodruff, G., Murase, K., Kitazume, H., Chai, C., Akagi, A., Panda, O., Ke, H., Schroeder, F., Wang, J., Berriman, M., Sternberg, P., Sugimoto, A.,

Kikuchi, T., 2018. Biology and genome of a newly discovered sibling species of *Caenorhabditis elegans*. Nat. Commun. 9, 3216.
Kern, C.C., Townsend, S., Salzmann, A., Rendell, N., Taylor, G., Comisel, R.M., Foukas, L., Bähler, J., Gems, D., 2020. *C. elegans* provide milk for their young. BioRxiv doi.org/10.1101/2020.11.15.380253.
Ludewig, A.H., Gimond, C., Judkins, J.C., Thornton, S., Pulido, D.C., Micikas, R.J., Döring, F., Antebi, A., Braendle, C., Schroeder, F.C., 2017. Larval crowding accelerates *C. elegans* development and reduces lifespan. PLOS Genet. 13, e1006717.
Maures, T., Booth, L., Benayoun, B., Izrayelit, Y., Schroeder, F., Brunet, A., 2014. Males shorten the life span of *C. elegans* hermaphrodites via secreted compounds. Science 343, 541-544.
Riesen, M., Feyst, I., Rattanavirotkul, N., Ezcurra, M., Tullet, J., Papatheodorou, I., Ziehm, M., Au, C., Gilliat, A., Hellberg, J., Thornton, J., Gems, D., 2014. MDL-1, a growth- and tumor-suppressor, slows aging and prevents germline hyperplasia and hypertrophy in *C. elegans*. Aging 6, 98-117.
Shaham, S., 2006. WormBook: Methods in Cell Biology, in: The *C. elegans* Research Community, (Ed.), WormBook.
Sharrock, W., 1983. Yolk proteins of *Caenorhabditis elegans*. Dev. Biol. 96, 182-188.
Shi, C., Murphy, C.T., 2014. Mating induces shrinking and death in *Caenorhabditis* mothers. Science 343, 536-540.
Sornda, T., Ezcurra, M., Kern, C., Galimov, E.R., Au, C., de la Guardia, Y., Gems, D., 2019. Production of YP170 vitellogenins promotes intestinal senescence in *C. elegans*. J. Gerontol. A 74, 1180-1188.
Wang, H., Zhao, Y., Ezcurra, M., Benedetto, A., Gilliat, A., Hellberg, J., Ren, Z., Athigapanich, T., Girstmair, J., Telford, M.J., Dolphin, C.T., Zhang, Z., Gems, D., 2018. A parthenogenetic quasi-program causes teratoma-like tumors during aging in wild-type *C. elegans*. NPJ Aging Mech. Disease 4, 6.
Wei, J.-Z., Hale, K., Carta, L., Platzer, E., Wong, C., Fang, S.-C., Aroian, R.V., 2003. *Bacillus thuringiensis* crystal proteins that target nematodes. Proc. Natl. Acad. Sci. U S A 100, 2760-2765.
Zhao, Y., Gilliat, A.F., Ziehm, M., Turmaine, M., Wang, H., Ezcurra, M., Yang, C., Phillips, G., McBay, D., Zhang, W.B., Partridge, L., Pincus, Z., Gems, D., 2017. Two forms of death in aging *Caenorhabditis elegans*. Nat. Commun. 8, 15458.
Zhao, Y., Wang, H., Poole, R.J., Gems, D., 2019. A *fln-2* mutation affects lethal pathology and lifespan in *C. elegans*. Nat. Commun. 10, 5087.

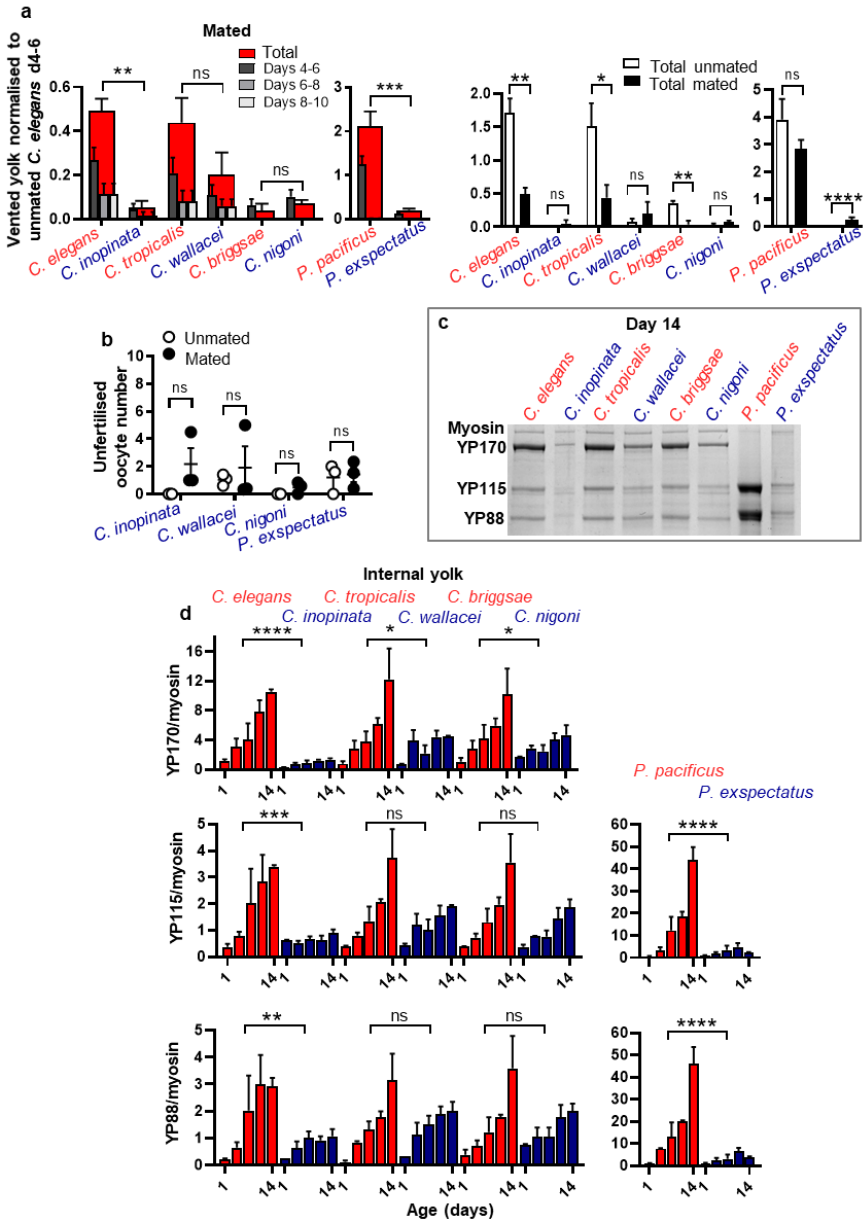

**Extended Data Figure 1 | Greater vitellogenin accumulation and venting in hermaphrodites than females. a**, Mating reduces venting in hermaphrodites and has little effect on venting by females. Vented YP170 in mated *Caenorhabditis* species and YP115 in mated *Pristionchus* species. Data normalised to unmated *C. elegans* peak venting period d4-6 (c.f. Fig.1b). Mating reduces levels of venting in hermaphrodites, which may reflect vitellogenin uptake by the additional oocytes that become eggs due to sperm from males. **b**, Mating does not significantly increase unfertilised oocyte production in females. **a**,**b**, One-way ANOVA (Bonferroni correction) and one-sample t-test (two-tailed) used based on the number of samples being compared. **c**,**d**, Greater internal levels of YP170 in *Caenorhabditis* hermaphrodites, and YP115/YP88 in *P. pacificus* hermaphrodites (*Pristionchus* lack YP170). YP bands normalised to myosin to adjust for species differences in body size. ANCOVA. **a**,**c**,**d**, Protein gel electrophoresis data with colloidal coomassie blue staining. Mean ± s.e.m. of 3 trials displayed. * $p$ <0.05, ** $p$ <0.01, *** $p$ <0.0001, **** $p$ <0.00001.

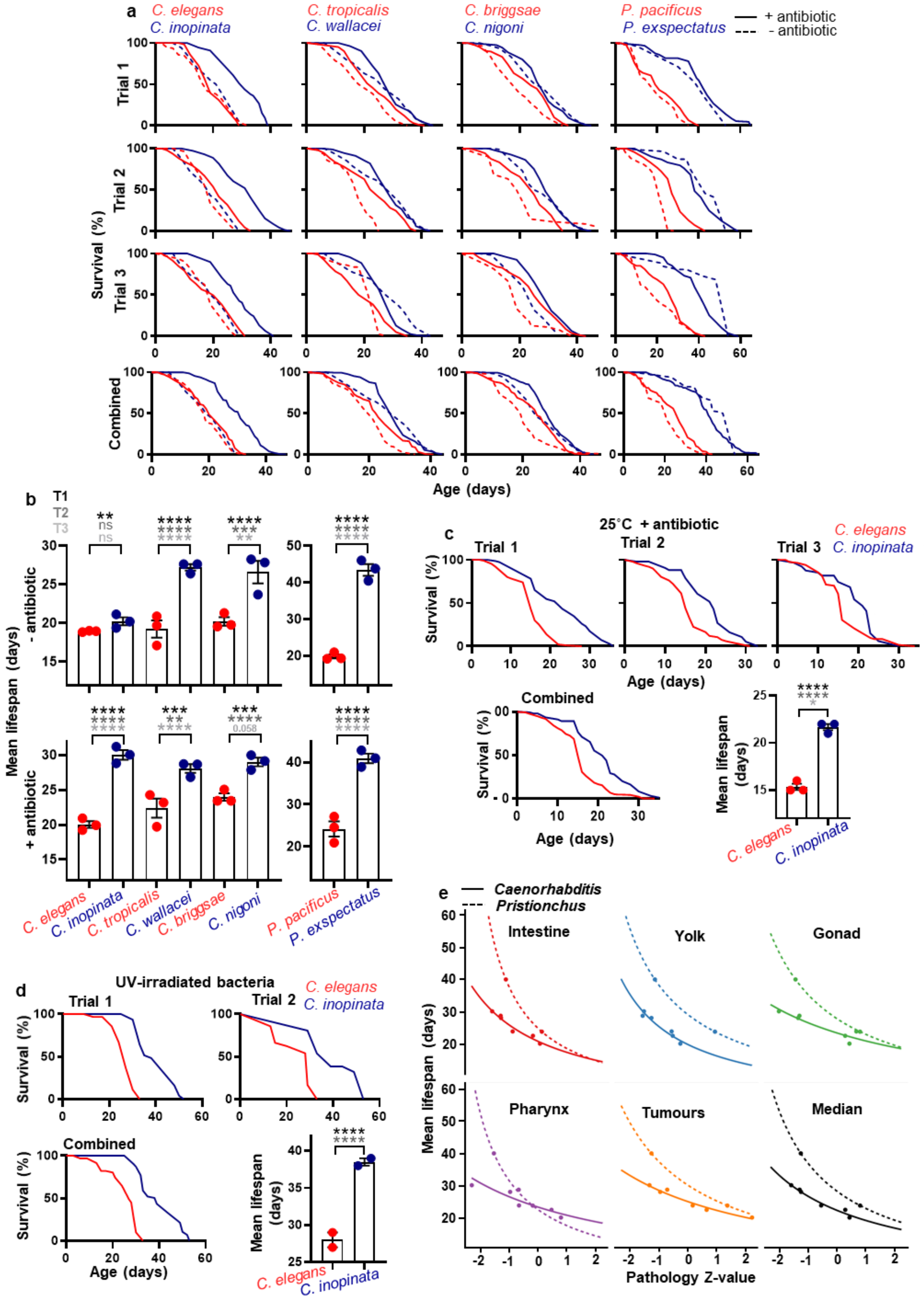


**Extended Data Figure 2 | More severe senescent pathology and shorter lifespan in hermaphroditic species. a**,**b**, Survival curves from individual and combined trials of sibling species. In the presence of antibiotics, females are longer lived in all cases. **c**, *C. inopinata* is also longer lived than *C. elegans* at 25˚C (with carbenicillin), the optimal culture temperature for the former species. **d**, *C. inopinata* is also longer-lived than *C. elegans* where both are maintained on UV-irradiated *E. coli*. **e**, Linear regression comparing each individual pathology and median of all pathologies to lifespan. The Z-scores for all pathologies were found to be statistically significant, and the median pathology

Z-score was found to perform better than the individual pathologies. This suggests that lifespan is a function of overall pathology level. X axis, pathology Z-score calculated from pathology measurements through time (c.f. Extended Data Fig. 4a). Y axis, mean lifespan. A line of best fit is drawn for *Caenorhabditis* species and based on this a hypothetical line is drawn for the *Pristionchus* species. * $p$ <0.05, ** $p$ <0.01, *** $p$ <0.0001, **** $p$ <0.00001, log rank test for pairs of individual trials. For raw lifespan data see Supplementary File 1.

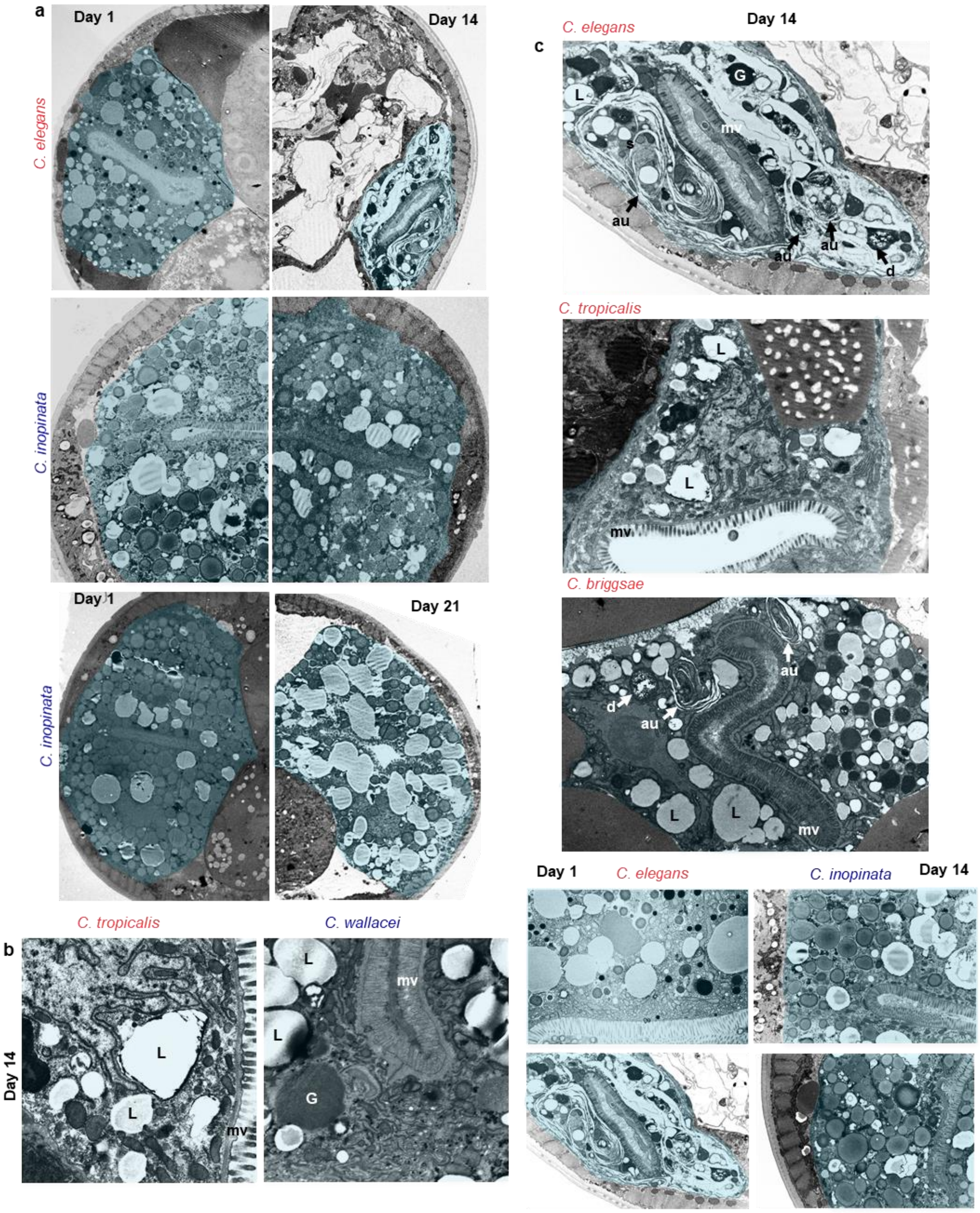


**Extended Data Figure 3 | Degenerative changes in ultrastructure in hermaphrodites undergoing reproductive death. a**, Low magnification representative TEM showing intestinal atrophy in *C. elegans* with age and little change in intestinal size in *C. inopinata* even by d21. Halves of nematode sibling species pairs are taken from approximately the same region of the animal (between the posterior end of the gonad and anus) to avoid the distorting effects of uterine tumours. Scale 5 µm (5,000x). **b**, *C. tropicalis* and *C. wallacei* on d14 showing greater ultrastructural degeneration in the former (c.f. Fig. 2d). Scale 1 µm (20,000x). **c**, Hermaproditic *Caenorhabditis* species showing intestinal degeneration on d14 and comparison to *C. inopinata*. Scale 1 µm (10,000x). Blue, intestine. mv: microvilli; au: autophagosomes; note characteristic multilamellar structure in some cases; L: lipid droplets; G: yolk granules or gut granules (tentative interpretation; gut granules have yet to be clearly identified in TEM studies; D. Hall personal communication).

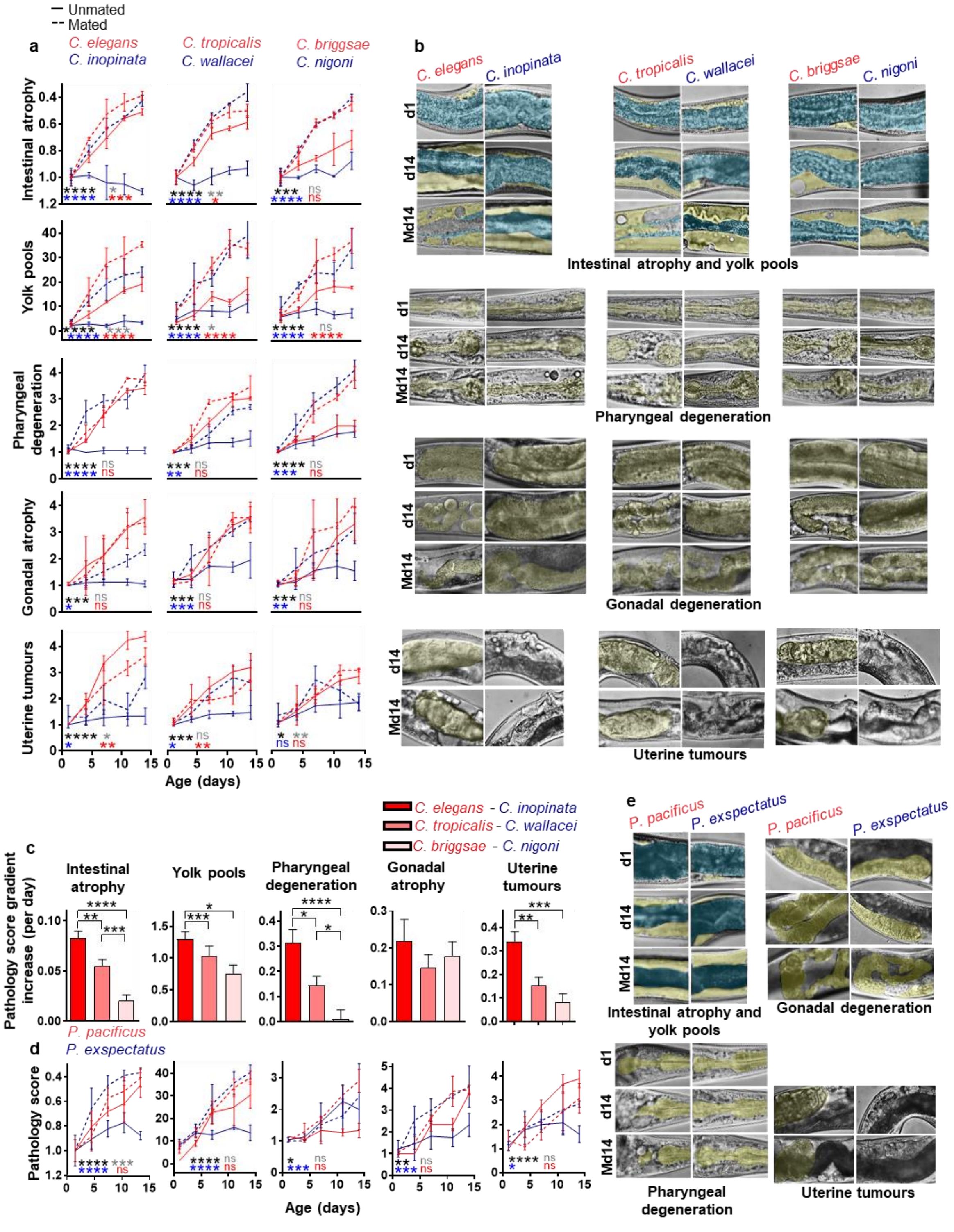


**Extended Data Figure 4 | Reproductive death is constitutive in hermaphrodites and facultative (mating-induced) in females. a**,**d**, Severe pathology progression in unmated hermaphrodites and mated animals but not virgin females. Tests for statistical significance: black, hermaphrodites vs females; red: mated vs unmated hermaphrodites; blue: mated vs unmated females; grey: mated hermaphrodites vs mated females. Cumulative Link Model with gamma link function for pharyngeal degeneration, gonadal degeneration and uterine tumours which are scored ordinals. ANCOVA with score normalised to d1 to determine change in percentage of intestinal mass, and ANCOVA with no normalization for yolk pools. Mean ± s.e.m. of 3 trials displayed (n=10 per time point per trial). **b**,**e**, Representative Nomarski microscopy images taken for the purpose of scoring/measuring pathologies. Note position of gonad around the intestine is different in *Pristionchus* species, with the full gonad arm visible in the same plane only after intestinal atrophy. Yellow: yolk pools, gonad and tumours;

blue: intestine. **c**, Bar graphs showing difference in gradient between species pairs, defining the following gradient in difference in severity for most pathologies: *C. elegans* - *C. inopinata* > *C. tropicalis* - *C. wallacei* > *C. briggsae* - *C. nigoni*. Subtraction rather than normalisation is used as gradients are already relative values. * $p$ <0.05, ** $p$ <0.01, *** $p$ <0.0001, **** $p$ <0.00001.

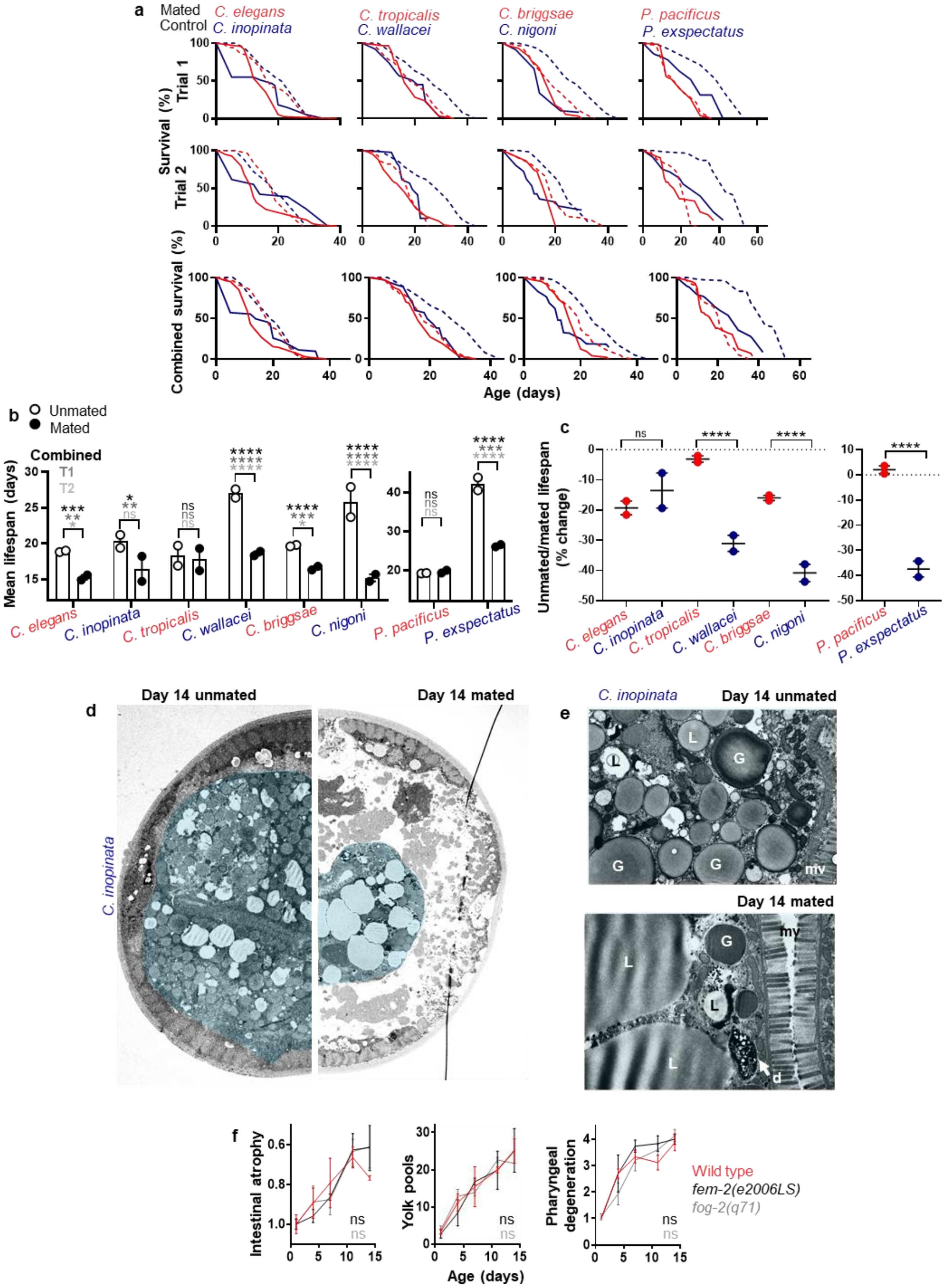


**Extended Data Figure 5 | Mating induces reproductive death and reduces lifespan in females.** **a**,**b**,**c**, Mating in females significantly reduces lifespan, as previously shown for *C. elegans* (Gems and Riddle, 1996; Maures et al., 2014; Shi and Murphy, 2014). **b**, Log rank between mated and unmated animals. **c**, Cox proportional hazard showing greater effect post mating in most females; the lack of a greater reduction in *C. inopinata* than *C. elegans* could reflect the greater susceptibility of the former to bacterial infection. **b**,**c**, Mean ± s.e.m. of 2 trials displayed. For raw lifespan data see Supplementary

File 1. **d**, Representative TEM cross-section showing a reduction in intestinal size in *C. inopinata* upon mating. Scale 5 µm (5,000x). **e**, Representative high magnification TEM images showing degenerative ultrastructural changes in the intestine in *C. inopinata* upon mating, with loss of ground structure similar to unmated *C. elegans,* and in contrast to unmated *C. inopinata,* observed (c.f. Fig. 1d). Scale 1 µm (20,000x). **f**, Reproductive death in *C. elegans* is not a consequence of the presence of self-sperm. Female mutant *C. elegans* show no significant difference in intestinal atrophy, yolk pool accumulation or pharyngeal atrophy (25˚C); it was previously shown that spermlessness accelerates uterine tumour growth which enhances gonad atrophy (Wang et al., 2018). * $p$ <0.05, ** $p$ <0.01, *** $p$ <0.0001, **** $p$ <0.00001.

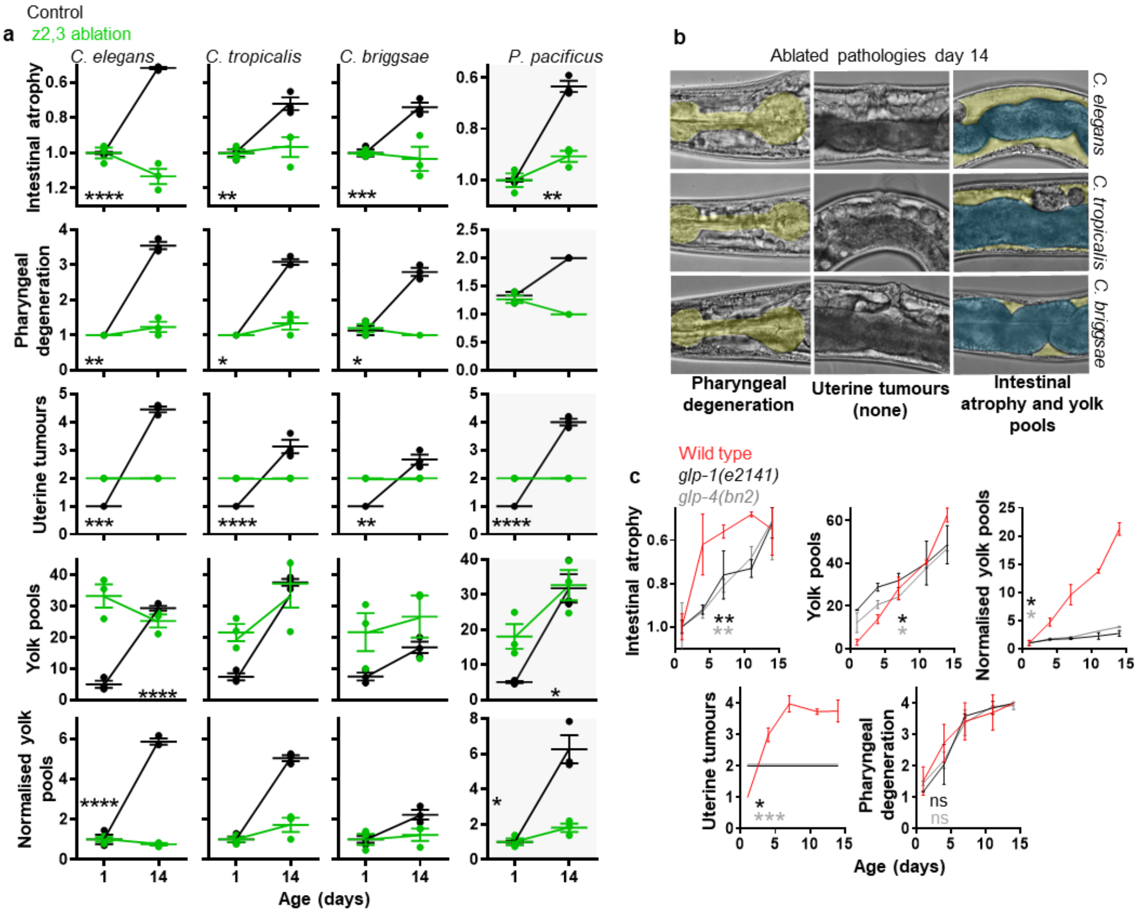


**Extended Data Figure 6 | Germline ablation suppresses reproductive death associated pathologies in hermaphrodites. a**, Pathology progression in hermaphrodites is suppressed following z2,3 ablation. **b**, Representative Nomarski microscopy images taken for the purpose of scoring pathologies. **c**, Pathology progression in *C. elegans* germline-deficient mutants showing a reduction in most RD-associated pathologies (25˚C). Note that suppression of germline development is not complete in these mutants, in contrast to laser ablated animals, which may account for the weaker suppression of RD-associated pathologies. Mean ± s.e.m. of 3 trials displayed. * $p$ <0.05, ** $p$ <0.01, *** $p$ <0.0001, **** $p$ <0.00001.

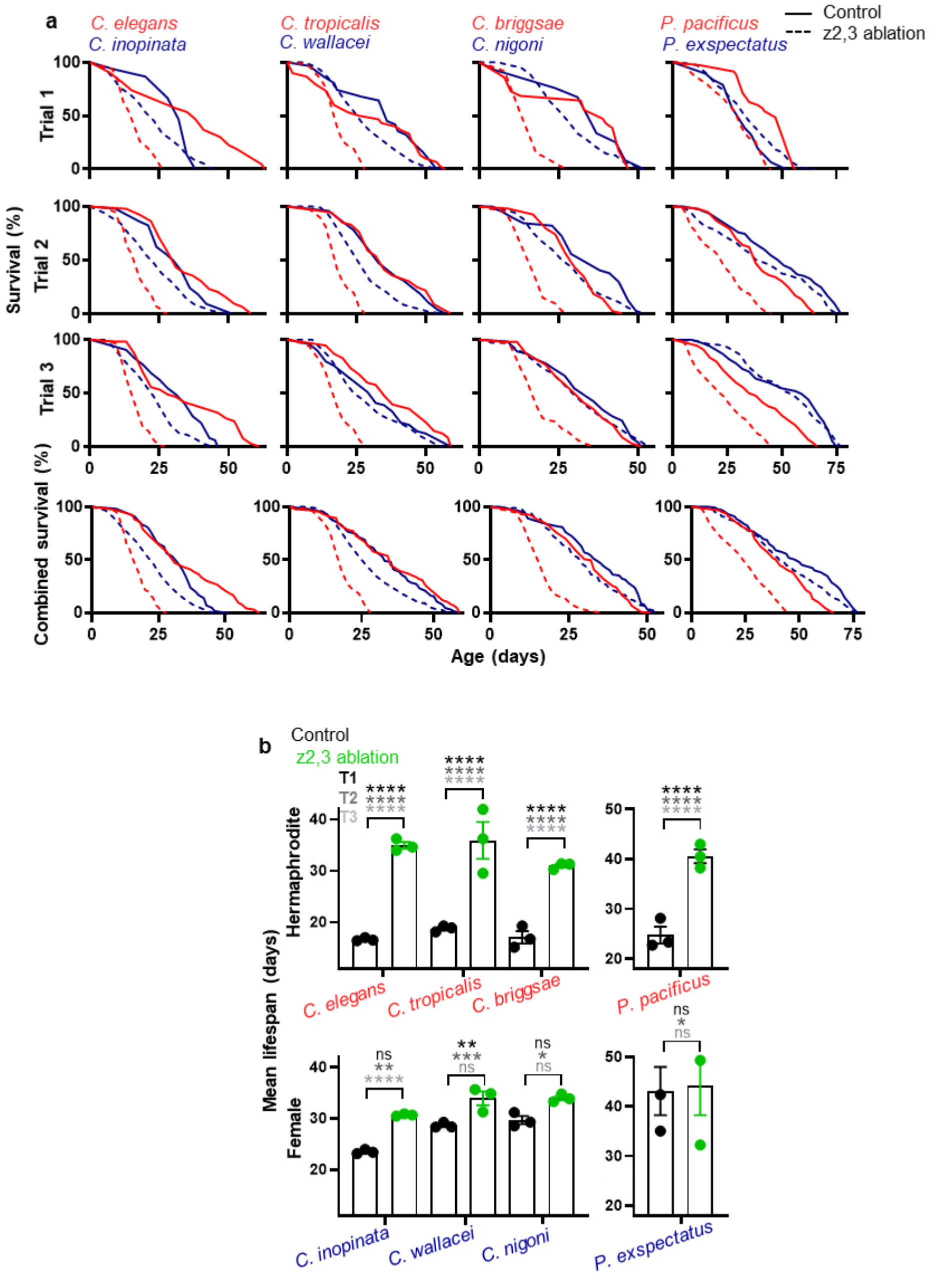


**Extended Data Figure 7 | Germline ablation greatly extends lifespan in hermaphrodites but not females. a**,**b**, Combined lifespans and individual trials of sibling species showing lifespan extension post z2,3 ablation only has a large effect in *Caenorhabditis* and *Pristionchus* hermaphrodites. Log rank tests for individual trials. Mean ± s.e.m. of 3 trials displayed. * $p$ <0.05, ** $p$ <0.01, *** $p$ <0.0001, **** $p$ <0.00001. For details on statistics and raw lifespan data see Extended data Table 2 and Supplementary File 1, respectively.

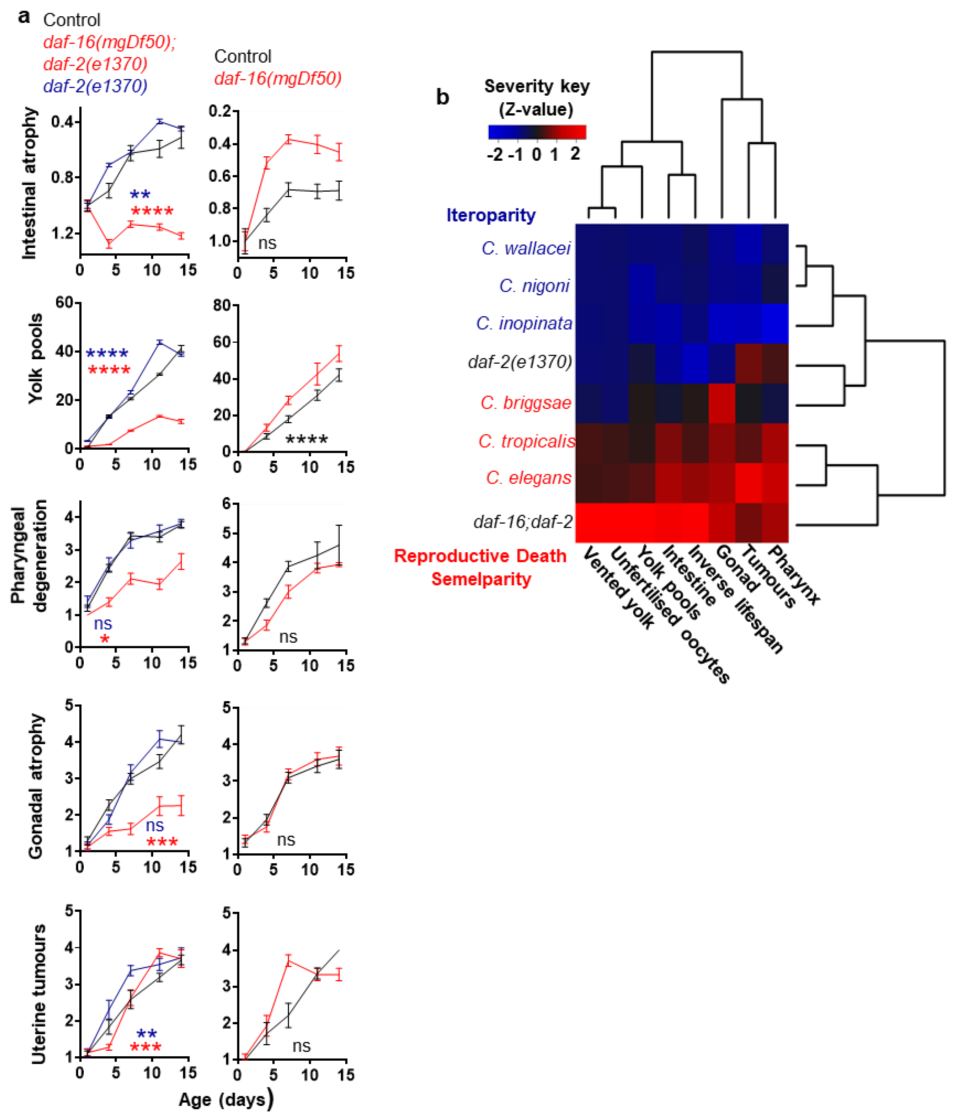


**Extended Data Figure 8 | Reproductive death is promoted by insulin/IGF-1 signalling (IIS). a**, Pathology progression is suppressed in *daf-2(e1370)* mutants, and this is *daf-16* dependant. Mean ± s.e.m. of 3 trials displayed. * $p$ <0.05, ** $p$ <0.01, *** $p$ <0.0001, **** $p$ <0.00001. **b**, Lifespan, reproductive death associated pathologies and yolk-lactation (yolk venting and unfertilised oocyte production) follow a continuum with *daf-2* behaving like unmated females and *daf-16; daf-2* mutants like unmated hermaphrodites. Heat map comparing differences in pathology progression, lifespan, yolk venting and oocyte production across species and treatments by transforming the calculated gradients of pathology progression into Z-scores. Hierarchical clustering based on pair-wise Euclidean distances was used to cluster pathologies and species/treatments according to similarity. Sources: Mutant yolk and oocyte venting data: Kern et al. (2020); mutant lifespan data: Bansal et al. (2014).

| antibiotic and control 20℃ | | Number | | Lifespan (days) | | %change treatment | %change ♀ vs ⚥ | Logrank | Cox PH Prob>ChiSq |
|---|---|---|---|---|---|---|---|---|---|
| | | death | censored | mean | median | | | | |
| | C | 219 | 27 | 20.14 | 20.00 | 6.08 | 49.92 | ⚥T vs ⚥ 0.0049 | <.0001 |
| *C. elegans* antibiotic | 1 | 55 | 9 | 19.93 | 19.00 | 5.75 | 50.82 | ⚥T vs ⚥ 0.8516 | <.0001 |
| | 2 | 92 | 5 | 20.92 | 22.00 | 10.34 | 49.12 | ⚥T vs ⚥ 0.0012 | <.0001 |
| | 3 | 72 | 13 | 19.25 | 20.00 | 1.22 | 49.75 | ⚥T vs ⚥ 0.1069 | 0.0087 |
| | C | 216 | 31 | 18.99 | 19.00 | | 6.79 | ⚥ vs ♀ 0.0057 | |
| *C. elegans* control | 1 | 112 | 7 | 18.85 | 18.00 | | 12.22 | ⚥ vs ♀ 0.1287 | |
| | 2 | 54 | 11 | 18.96 | 19.00 | | 2.26 | ⚥ vs ♀ 0.1192 | |
| | 3 | 50 | 13 | 19.02 | 19.00 | | 6.25 | ⚥ vs ♀ 0.0491 | |
| | C | 172 | 111 | 30.20 | 31.00 | 48.92 | | ♀T vs ♀ < 0.0001 | |
| *C. inopinata* antibiotic | 1 | 51 | 39 | 30.06 | 31.00 | 42.12 | | ♀T vs ♀ < 0.0001 | |
| | 2 | 71 | 9 | 31.20 | 33.00 | 60.89 | | ♀T vs ♀ < 0.0001 | |
| | 3 | 50 | 63 | 28.83 | 29.00 | 42.65 | | ♀T vs ♀ < 0.0001 | |
| | C | 130 | 96 | 20.28 | 20.00 | | | ⚥T vs ♀T < 0.0001 | |
| *C. inopinata* control | 1 | 41 | 32 | 21.15 | 24.00 | | | ⚥T vs ♀T < 0.0001 | |
| | 2 | 43 | 32 | 19.39 | 19.00 | | | ⚥T vs ♀T < 0.0001 | |
| | 3 | 46 | 32 | 20.21 | 21.00 | | | ⚥T vs ♀T < 0.0001 | |
| | C | 264 | 22 | 22.55 | 23.00 | 15.71 | 24.75 | ⚥T vs ⚥ < 0.0001 | <.0001 |
| *C. tropicalis* antibiotic | 1 | 95 | 10 | 24.28 | 23.00 | 23.20 | 18.34 | ⚥T vs ⚥ 0.0003 | 0.0559 |
| | 2 | 91 | 5 | 23.17 | 25.00 | 35.46 | 23.73 | ⚥T vs ⚥ < 0.0001 | <.0001 |
| | 3 | 78 | 7 | 19.71 | 23.00 | -5.72 | 36.14 | ⚥T vs ⚥ 0.4462 | 0.0226 |
| | C | 162 | 43 | 19.49 | 21.00 | | 39.54 | ⚥ vs ♀ < 0.0001 | |
| *C. tropicalis* control | 1 | 78 | 7 | 19.71 | 23.00 | | 33.97 | ⚥ vs ♀ < 0.0001 | |
| | 2 | 44 | 16 | 17.11 | 18.00 | | 61.59 | ⚥ vs ♀ < 0.0001 | |
| | 3 | 40 | 20 | 20.90 | 23.00 | | 32.19 | ⚥ vs ♀ < 0.0001 | |
| | C | 230 | 72 | 28.13 | 27.00 | 3.44 | | ♀T vs ♀ 0.1461 | |
| *C. wallacei* antibiotic | 1 | 82 | 25 | 28.73 | 28.00 | 8.82 | | ♀T vs ♀ 0.6742 | |
| | 2 | 78 | 22 | 28.67 | 29.00 | 3.72 | | ♀T vs ♀ 0.7144 | |
| | 3 | 70 | 25 | 26.83 | 27.00 | -2.91 | | ♀T vs ♀ 0.0038 | |
| | C | 240 | 49 | 27.19 | 27.00 | | | ⚥T vs ♀T < 0.0001 | |
| *C. wallacei* control | 1 | 87 | 10 | 26.40 | 27.00 | | | ⚥T vs ♀T 0.0006 | |
| | 2 | 77 | 20 | 27.64 | 32.00 | | | ⚥T vs ♀T 0.0010 | |
| | 3 | 76 | 19 | 27.63 | 32.00 | | | ⚥T vs ♀T < 0.0001 | |
| | C | 273 | 29 | 23.91 | 25.00 | 20.33 | 20.52 | ⚥T vs ⚥ 0.0049 | 0.0165 |
| *C. briggsae* antibiotic | 1 | 107 | 9 | 23.44 | 24.00 | 18.53 | 18.92 | ⚥T vs ⚥ 0.0007 | 0.0056 |
| | 2 | 86 | 6 | 23.43 | 25.00 | 9.88 | 28.01 | ⚥T vs ⚥ 0.1113 | 0.8878 |
| | 3 | 80 | 14 | 25.07 | 25.00 | 28.06 | 16.33 | ⚥T vs ⚥ 0.0002 | 0.629 |
| | C | 190 | 34 | 19.87 | 20.00 | | 35.53 | ⚥ vs ♀ < 0.0001 | |
| *C. briggsae* control | 1 | 96 | 20 | 19.78 | 24.00 | | 40.80 | ⚥ vs ♀ < 0.0001 | |
| | 2 | 40 | 14 | 21.32 | 19.00 | | 32.56 | ⚥ vs ♀ 0.0023 | |
| | 3 | 54 | 0 | 19.57 | 19.00 | | 21.02 | ⚥ vs ♀ 0.0165 | |
| | C | 268 | 49 | 28.82 | 29.00 | 7.00 | | ♀T vs ♀ 0.228 | |
| *C. nigoni* antibiotic | 1 | 116 | 10 | 27.87 | 28.00 | 0.11 | | ♀T vs ♀ 0.5663 | |
| | 2 | 74 | 21 | 29.99 | 29.00 | 6.11 | | ♀T vs ♀ 0.4993 | |
| | 3 | 78 | 18 | 29.16 | 225.00 | 23.10 | | ♀T vs ♀ < 0.0001 | |
| | C | 227 | 46 | 26.93 | 25.00 | | | ⚥T vs ♀T < 0.0001 | |
| *C. nigoni* control | 1 | 90 | 11 | 27.84 | 28.00 | | | ⚥T vs ♀T 0.0004 | |
| | 2 | 76 | 21 | 28.26 | 28.00 | | | ⚥T vs ♀T < 0.0001 | |
| | 3 | 61 | 14 | 23.69 | 23.00 | | | ⚥T vs ♀T 0.0582 | |
| | C | 232 | 26 | 23.86 | 27.00 | 18.71 | 68.22 | ⚥T vs ⚥ < 0.0001 | <.0001 |
| *P. pacificus* antibiotic | 1 | 92 | 7 | 20.93 | 19.00 | 9.12 | 95.85 | ⚥T vs ⚥ 0.1019 | 0.7063 |
| | 2 | 68 | 6 | 27.12 | 27.00 | 40.36 | 43.54 | ⚥T vs ⚥ < 0.0001 | <.0001 |
| | 3 | 72 | 13 | 24.59 | 27.00 | 17.48 | 74.23 | ⚥T vs ⚥ 0.0674 | 0.018 |
| | C | 274 | 65 | 20.10 | 20.00 | | 120.37 | ⚥ vs ♀ < 0.0001 | |
| *P. pacificus* control | 1 | 93 | 12 | 19.18 | 20.00 | | 111.05 | ⚥ vs ♀ < 0.0001 | |
| | 2 | 79 | 41 | 19.32 | 21.00 | | 126.81 | ⚥ vs ♀ < 0.0001 | |
| | 3 | 102 | 12 | 20.93 | 20.00 | | 119.88 | ⚥ vs ♀ < 0.0001 | |
| | C | 271 | 46 | 40.14 | 41.00 | -9.38 | | ♀T vs ♀ 0.0111 | |
| *P. exspectatus* antibiotic | 1 | 110 | 11 | 40.99 | 42.00 | 1.26 | | ♀T vs ♀ 0.5208 | |
| | 2 | 81 | 11 | 38.92 | 38.00 | -11.17 | | ♀T vs ♀ 0.1355 | |
| | 3 | 105 | 24 | 42.85 | 45.00 | -6.91 | | ♀T vs ♀ 0.0017 | |
| | C | 73 | 58 | 44.29 | 53.00 | | | ⚥T vs ♀T < 0.0001 | |
| *P. exspectatus* control | 1 | 21 | 11 | 40.48 | 43.00 | | | ⚥T vs ♀T < 0.0001 | |
| | 2 | 29 | 44 | 43.82 | 43.00 | | | ⚥T vs ♀T < 0.0001 | |
| | 3 | 22 | 3 | 46.02 | 53.00 | | | ⚥T vs ♀T < 0.0001 | |

| antibiotic and control 25℃ | | Number | | Lifespan (days) | | %change ♀ vs ⚥ | Logrank | Cox PH Prob>ChiSq |
|---|---|---|---|---|---|---|---|---|
| | | death | censored | mean | median | | | |
| | C | 268 | 66 | 15.57 | 15.00 | -25.76 | ⚥T vs ♀T < 0.0001 | 0.0407 |
| *C. elegans* | 1 | 110 | 32 | 15.09 | 15.00 | -33.72 | ⚥T vs ♀T < 0.0001 | |
| | 2 | 83 | 22 | 15.30 | 15.00 | -27.64 | ⚥T vs ♀T < 0.0001 | |
| | 3 | 75 | 12 | 16.53 | 16.00 | -13.35 | ⚥T vs ♀T 0.0198 | |
| | C | 176 | 60 | 20.97 | 22.00 | | | |
| *C. inopinata* | 1 | 75 | 13 | 22.76 | 22.00 | | | |
| | 2 | 27 | 27 | 21.15 | 22.00 | | | |
| | 3 | 74 | 20 | 19.08 | 21.00 | | | |

| UV irradiated bacteria 20℃ | | Number | | Lifespan (days) | | %change ♀ vs ⚥ | Logrank | Cox PH Prob>ChiSq |
|---|---|---|---|---|---|---|---|---|
| | | death | censored | mean | median | | | |
| | C | 46 | 27 | 25.65 | 28.00 | -35.14 | ⚥T vs ♀T < 0.0001 | <.0001 |
| *C. elegans* | 1 | 24 | 18 | 27.54 | 27.00 | -31.01 | ⚥T vs ♀T < 0.0001 | |
| | 2 | 22 | 9 | 23.77 | 29.00 | -40.75 | ⚥T vs ♀T < 0.0001 | |
| | C | 43 | 27 | 39.55 | 35.00 | | | |
| *C. inopinata* | 1 | 30 | 20 | 39.92 | 38.00 | | | |
| | 2 | 13 | 7 | 40.12 | 33.00 | | | |

**Extended Data Table 1 | Female sibling species are longer lived than hermaphrodites when bacterial pathogenicity is accounted for.** C is combined data from all trials, and T is treatment (antibiotic carbenicillin vs control).

| Mated vs control 20℃ | | Number | | lifespan(days) | | %change treatment | %change ♀ vs ⚥ | Logrank | Cox PH Prob>ChiSq |
|---|---|---|---|---|---|---|---|---|---|
| | | death | censored | mean | median | | | | |
| *C. elegans* mated | C | 123 | 24 | 15.01 | 12.00 | -20.72 | 5.58 | ⚥T vs ⚥ 0.0004 | 0.8463 |
| | 1 | 41 | 8 | 15.62 | 16.00 | -17.14 | -5.89 | ⚥T vs ⚥ 0.0017 | 0.5995 |
| | 2 | 82 | 16 | 14.91 | 12.00 | -21.36 | 21.82 | ⚥T vs ⚥ 0.0194 | 0.1326 |
| *C. elegans* control | C | 166 | 18 | 18.93 | 18.00 | | 7.19 | ⚥ vs ♀ 0.0455 | |
| | 1 | 112 | 7 | 18.85 | 18.00 | | 12.22 | ⚥ vs ♀ 0.1287 | |
| | 2 | 54 | 11 | 18.96 | 19.00 | | 2.26 | ⚥ vs ♀ 0.1192 | |
| *C. inopinata* mated | C | 105 | 16 | 15.84 | 14.00 | -21.91 | | ♀T vs ♀ 0.0213 | |
| | 1 | 74 | 13 | 14.70 | 18.00 | -30.51 | | ♀T vs ♀ 0.0012 | |
| | 2 | 31 | 3 | 18.17 | 14.00 | -6.31 | | ♀T vs ♀ 0.6203 | |
| *C. inopinata* control | C | 84 | 64 | 20.29 | 19.00 | | | ⚥T vs ♀T 0.1954 | |
| | 1 | 41 | 32 | 21.15 | 24.00 | | | ⚥T vs ♀T 0.6610 | |
| | 2 | 43 | 32 | 19.39 | 19.00 | | | ⚥T vs ♀T 0.1117 | |
| *C. tropicalis* mated | C | 144 | 7 | 18.07 | 16.00 | -3.54 | 10.64 | ⚥T vs ⚥ 0.7607 | <.0001 |
| | 1 | 83 | 7 | 19.31 | 16.00 | -2.00 | -2.33 | ⚥T vs ⚥ 0.8952 | 0.0069 |
| | 2 | 61 | 0 | 16.20 | 16.00 | -4.12 | 13.16 | ⚥T vs ⚥ 0.9253 | 0.0017 |
| *C. tropicalis* control | C | 123 | 22 | 18.73 | 18.00 | | 44.13 | ⚥ vs ♀ < 0.0001 | |
| | 1 | 78 | 7 | 19.71 | 23.00 | | 33.97 | ⚥ vs ♀ < 0.0001 | |
| | 2 | 45 | 15 | 16.89 | 18.00 | | 63.63 | ⚥ vs ♀ < 0.0001 | |
| *C. wallacei* mated | C | 83 | 55 | 19.99 | 19.00 | -25.96 | | ♀T vs ♀ < 0.0001 | |
| | 1 | 30 | 21 | 18.86 | 21.00 | -28.56 | | ♀T vs ♀ 0.0048 | |
| | 2 | 53 | 34 | 18.33 | 19.00 | -33.69 | | ♀T vs ♀ < 0.0001 | |
| *C. wallacei* control | C | 164 | 30 | 27.00 | 27.00 | | | ⚥T vs ♀T 0.0441 | |
| | 1 | 87 | 10 | 26.40 | 27.00 | | | ⚥T vs ♀T 0.5469 | |
| | 2 | 77 | 20 | 27.64 | 32.00 | | | ⚥T vs ♀T 0.0185 | |
| *C. briggsae* mated | C | 180 | 1 | 16.62 | 16.00 | -15.34 | -12.04 | ⚥T vs ⚥ < 0.0001 | <.0001 |
| | 1 | 123 | 0 | 16.78 | 16.00 | -15.14 | -6.17 | ⚥T vs ⚥ 0.0004 | <.0001 |
| | 2 | 57 | 1 | 16.27 | 16.00 | -16.90 | -9.41 | ⚥T vs ⚥ 0.0159 | 0.1378 |
| *C. briggsae* control | C | 150 | 20 | 19.63 | 20.00 | | 33.53 | ⚥ vs ♀ < 0.0001 | |
| | 1 | 96 | 20 | 19.78 | 20.00 | | 40.86 | ⚥ vs ♀ < 0.0001 | |
| | 2 | 54 | 0 | 19.57 | 19.00 | | 21.02 | ⚥ vs ♀ 0.0165 | |
| *C. nigoni* mated | C | 161 | 66 | 14.62 | 12.00 | -44.23 | | ♀T vs ♀ < 0.0001 | |
| | 1 | 43 | 4 | 15.74 | 14.00 | -43.48 | | ♀T vs ♀ < 0.0001 | |
| | 2 | 118 | 62 | 14.74 | 12.00 | -37.80 | | ♀T vs ♀ < 0.0001 | |
| *C. nigoni* control | C | 151 | 26 | 26.22 | 25.00 | | | ⚥T vs ♀T 0.0746 | |
| | 1 | 90 | 12 | 27.86 | 28.00 | | | ⚥T vs ♀T 0.1812 | |
| | 2 | 61 | 14 | 23.69 | 23.00 | | | ⚥T vs ♀T 0.1748 | |
| *P. pacificus* mated | C | 132 | 50 | 19.68 | 19.00 | 0.99 | 33.85 | ⚥T vs ⚥ 0.2286 | <.0001 |
| | 1 | 59 | 18 | 19.30 | 19.00 | 0.62 | 37.82 | ⚥T vs ⚥ 0.8972 | 0.0027 |
| | 2 | 73 | 32 | 19.95 | 19.00 | 3.27 | 30.31 | ⚥T vs ⚥ 0.5063 | <.0001 |
| *P. pacificus* control | C | 172 | 53 | 19.48 | 21.00 | | 120.08 | ⚥ vs ♀ < 0.0001 | |
| | 1 | 93 | 12 | 19.18 | 20.00 | | 111.05 | ⚥ vs ♀ < 0.0001 | |
| | 2 | 79 | 41 | 19.32 | 21.00 | | 126.81 | ⚥ vs ♀ < 0.0001 | |
| *P. exspectatus* mated | C | 82 | 32 | 26.34 | 30.00 | -38.58 | | ♀T vs ♀ < 0.0001 | |
| | 1 | 39 | 16 | 26.60 | 30.00 | -34.29 | | ♀T vs ♀ 0.0019 | |
| | 2 | 43 | 16 | 26.00 | 30.00 | -40.67 | | ♀T vs ♀ < 0.0001 | |
| *P. exspectatus* control | C | 50 | 55 | 42.88 | 43.00 | | | ⚥T vs ♀T < 0.0001 | |
| | 1 | 21 | 11 | 40.48 | 43.00 | | | ⚥T vs ♀T < 0.0001 | |
| | 2 | 29 | 44 | 43.82 | 43.00 | | | ⚥T vs ♀T 0.0107 | |

**Extended Data Table 2 | Comparison of hermaphrodite and female sibling species lifespans following mating.** C is combined data from all trials, and T is treatment (mated or unmated).

| z2,3, ablation 20℃ | | Number | | lifespan(days) | | %change treatment | %change ♀ vs ⚥ | Logrank | Cox PH Prob>ChiSq |
|---|---|---|---|---|---|---|---|---|---|
| | | death | censored | mean | median | | | | |
| | C | 116 | 22 | 34.71 | 31.00 | 107.89 | -11.83 | ⚥T vs ⚥ < 0.0001 | <.0001 |
| ***C. elegans*** | 1 | 28 | 4 | 36.23 | 38.00 | 120.18 | -15.35 | ⚥T vs ⚥ < 0.0001 | <.0001 |
| **ablated** | 2 | 43 | 9 | 34.65 | 31.00 | 103.14 | -10.76 | ⚥T vs ⚥ < 0.0001 | <.0001 |
| | 3 | 45 | 9 | 33.99 | 28.00 | 105.28 | -10.34 | ⚥T vs ⚥ < 0.0001 | <.0001 |
| | C | 282 | 18 | 16.70 | 18.00 | | 40.63 | ⚥ vs ♀ < 0.0001 | |
| ***C. elegans*** | 1 | 100 | 1 | 16.45 | 18.00 | | 40.56 | ⚥ vs ♀ < 0.0001 | |
| **control** | 2 | 96 | 3 | 17.06 | 18.00 | | 40.60 | ⚥ vs ♀ < 0.0001 | |
| | 3 | 86 | 14 | 16.56 | 15.00 | | 41.16 | ⚥ vs ♀ < 0.0001 | |
| | C | 110 | 14 | 30.61 | 32.00 | 30.34 | | ♀T vs ♀ < 0.0001 | |
| ***C. inopinata*** | 1 | 15 | 0 | 30.67 | 32.00 | 32.60 | | ♀T vs ♀ 0.1410 | |
| **ablated** | 2 | 45 | 12 | 30.92 | 30.00 | 28.94 | | ♀T vs ♀ 0.0011 | |
| | 3 | 50 | 2 | 30.48 | 31.00 | 30.38 | | ♀T vs ♀ < 0.0001 | |
| | C | 383 | 37 | 23.48 | 23.00 | | | ⚥T vs ♀T 0.0003 | |
| ***C. inopinata*** | 1 | 119 | 16 | 23.13 | 23.00 | | | ⚥T vs ♀T 0.0202 | |
| **control** | 2 | 119 | 6 | 23.98 | 23.00 | | | ⚥T vs ♀T 0.0436 | |
| | 3 | 145 | 15 | 23.38 | 23.00 | | | ⚥T vs ♀T 0.0242 | |
| | C | 118 | 20 | 35.21 | 35.00 | 87.21 | -3.92 | ⚥T vs ⚥ < 0.0001 | <.0001 |
| ***C. tropicalis*** | 1 | 30 | 0 | 29.53 | 29.50 | 55.97 | 18.07 | ⚥T vs ⚥ < 0.0001 | 0.0027 |
| **ablated** | 2 | 38 | 16 | 36.26 | 35.00 | 87.72 | -1.61 | ⚥T vs ⚥ < 0.0001 | <.0001 |
| | 3 | 34 | 20 | 41.94 | 47.00 | 130.90 | -25.45 | ⚥T vs ⚥ < 0.0001 | <.0001 |
| | C | 213 | 41 | 18.81 | 18.00 | | 52.46 | ⚥ vs ♀ < 0.0001 | |
| ***C. tropicalis*** | 1 | 70 | 14 | 18.93 | 18.00 | | 49.69 | ⚥ vs ♀ < 0.0001 | |
| **control** | 2 | 68 | 21 | 19.32 | 18.00 | | 46.77 | ⚥ vs ♀ < 0.0001 | |
| | 3 | 75 | 6 | 18.16 | 18.00 | | 61.10 | ⚥ vs ♀ < 0.0001 | |
| | C | 129 | 1 | 33.83 | 33.00 | 17.98 | | ♀T vs ♀ < 0.0001 | |
| ***C. wallacei*** | 1 | 31 | 0 | 34.87 | 36.00 | 23.03 | | ♀T vs ♀ 0.0057 | |
| **ablated** | 2 | 49 | 1 | 35.68 | 33.00 | 25.83 | | ♀T vs ♀ 0.0002 | |
| | 3 | 49 | 0 | 31.27 | 32.00 | 6.86 | | ♀T vs ♀ 0.3265 | |
| | C | 386 | 30 | 28.67 | 27.00 | | | ⚥T vs ♀T 0.1631 | |
| ***C. wallacei*** | 1 | 137 | 15 | 28.34 | 27.00 | | | ⚥T vs ♀T 0.7572 | |
| **control** | 2 | 122 | 12 | 28.35 | 27.00 | | | ⚥T vs ♀T 0.7531 | |
| | 3 | 127 | 3 | 29.26 | 25.50 | | | ⚥T vs ♀T 0.0801 | |
| | C | 110 | 22 | 30.90 | 32.00 | 80.13 | 8.73 | ⚥T vs ⚥ < 0.0001 | <.0001 |
| **C. briggsae** | 1 | 24 | 5 | 31.33 | 39.00 | 106.39 | 11.06 | ⚥T vs ⚥ < 0.0001 | 0.0001 |
| **ablated** | 2 | 49 | 5 | 30.26 | 30.00 | 80.64 | 11.99 | ⚥T vs ⚥ < 0.0001 | <.0001 |
| | 3 | 37 | 12 | 31.42 | 30.00 | 62.70 | 5.92 | ⚥T vs ⚥ < 0.0001 | <.0001 |
| | C | 264 | 12 | 17.15 | 18.00 | | 73.08 | ⚥ vs ♀ < 0.0001 | |
| ***C. briggsae*** | 1 | 88 | 0 | 15.18 | 15.00 | | 92.91 | ⚥ vs ♀ < 0.0001 | |
| **control** | 2 | 82 | 8 | 16.75 | 18.00 | | 70.59 | ⚥ vs ♀ < 0.0001 | |
| | 3 | 94 | 4 | 19.31 | 18.00 | | 61.50 | ⚥ vs ♀ < 0.0001 | |
| | C | 101 | 16 | 33.59 | 34.00 | 13.15 | | ♀T vs ♀ 0.0152 | |
| ***C. nigoni* ablated** | 1 | 12 | 2 | 34.80 | 37.00 | 18.82 | | ♀T vs ♀ 0.1412 | |
| | 2 | 38 | 9 | 33.88 | 35.00 | 18.58 | | ♀T vs ♀ 0.0140 | |
| | 3 | 51 | 5 | 33.28 | 33.00 | 6.70 | | ♀T vs ♀ 0.6898 | |
| | C | 371 | 49 | 29.69 | 29.00 | | | ⚥T vs ♀T 0.0061 | |
| ***C. nigoni* control** | 1 | 120 | 20 | 29.29 | 28.00 | | | ⚥T vs ♀T 0.9253 | |
| | 2 | 123 | 12 | 28.57 | 29.00 | | | ⚥T vs ♀T 0.0050 | |
| | 3 | 128 | 17 | 31.19 | 32.00 | | | ⚥T vs ♀T 0.2840 | |
| | C | 138 | 11 | 40.23 | 41.00 | 64.28 | 20.50 | ⚥T vs ⚥ < 0.0001 | <.0001 |
| ***P. pacificus*** | 1 | 35 | 0 | 43.00 | 47.00 | 52.86 | -24.98 | ⚥T vs ⚥ < 0.0001 | <.0001 |
| **ablated** | 2 | 54 | 6 | 40.49 | 38.00 | 78.34 | 21.82 | ⚥T vs ⚥ < 0.0001 | 0.0095 |
| | 3 | 49 | 5 | 38.21 | 35.00 | 63.55 | 34.00 | ⚥T vs ⚥ < 0.0001 | <.0001 |
| | C | 236 | 83 | 24.49 | 25.00 | | 76.91 | ⚥ vs ♀ < 0.0001 | |
| ***P. pacificus*** | 1 | 53 | 54 | 28.13 | 31.00 | | 24.55 | ⚥ vs ♀ 0.0002 | |
| **control** | 2 | 90 | 18 | 22.71 | 23.00 | | 86.68 | ⚥ vs ♀ < 0.0001 | |
| | 3 | 93 | 11 | 23.36 | 23.00 | | 122.01 | ⚥ vs ♀ < 0.0001 | |
| | C | 169 | 12 | 48.47 | 50.00 | 11.90 | | ♀T vs ♀ 0.0020 | |
| ***P. exspectatus*** | 1 | 17 | 2 | 32.26 | 33.00 | -7.93 | | ♀T vs ♀ 0.1298 | |
| **ablated** | 2 | 74 | 3 | 49.33 | 51.00 | 16.38 | | ♀T vs ♀ 0.0081 | |
| | 3 | 78 | 7 | 51.20 | 59.00 | -1.28 | | ♀T vs ♀ 0.7380 | |
| | C | 367 | 111 | 43.32 | 41.00 | | | ⚥T vs ♀T < 0.0001 | |
| ***P. exspectatus*** | 1 | 125 | 10 | 35.04 | 35.00 | | | ⚥T vs ♀T 0.0002 | |
| **control** | 2 | 105 | 50 | 42.39 | 41.00 | | | ⚥T vs ♀T < 0.0001 | |
| | 3 | 137 | 51 | 51.87 | 55.00 | | | ⚥T vs ♀T < 0.0001 | |

**Extended Data Table 3 | Comparison of lifespans of hermaphrodite and female sibling species following germline ablation.** C is combined data from all trials, and T is treatment (z2,3 ablation vs control).
